# HDAC11 inhibition triggers bimodal thermogenic pathways to circumvent adipocyte catecholamine resistance

**DOI:** 10.1101/2023.03.29.534830

**Authors:** Emma L. Robinson, Rushita A. Bagchi, Jennifer L. Major, Bryan C. Bergman, Jennifer L. Madsuda, Timothy A. McKinsey

## Abstract

Stimulation of adipocyte β-adrenergic receptors (β-ARs) induces expression of uncoupling protein 1 (UCP1), promoting non-shivering thermogenesis. Association of β-ARs with a lysine myristoylated form of A-kinase anchoring protein 12 (AKAP12)/gravin-α is required for downstream signaling that culminates in UCP1 induction. Conversely, demyristoylation of gravin-α by histone deacetylase 11 (HDAC11) suppresses this pathway. Whether inhibition of HDAC11 in adipocytes is sufficient to drive UCP1 expression independently of β-ARs is not known. Here, we demonstrate that adipocyte-specific deletion of HDAC11 in mice leads to robust induction of UCP1 in adipose tissue (AT), resulting in increased body temperature. These effects are mimicked by treating mice *in vivo* or human AT *ex vivo* with an HDAC11-selective inhibitor, FT895. FT895 triggers biphasic, gravin-α myristoylation-dependent induction of UCP1 protein expression, with a non-canonical acute response that is post-transcriptional and independent of protein kinase A (PKA), and a delayed response requiring PKA activity and new *Ucp1* mRNA synthesis. Remarkably, HDAC11 inhibition promotes UCP1 expression even in models of adipocyte catecholamine resistance where β-AR signaling is blocked. These findings define cell autonomous, multi-modal roles for HDAC11 as a suppressor of thermogenesis, and highlight the potential of inhibiting HDAC11 to therapeutically alter AT phenotype independently of β-AR stimulation.

## Introduction

The prevalence of obesity in adults in the United States was recently estimated to be >40%, and this number is expected to rise in coming years given that ∼20% of children aged 2 – 19 years are already obese or overweight (1,2). Significant advancements in pharmacotherapy for obesity have recently been realized, as exemplified by glucagon-like peptide-1 receptor agonists, which reduce appetite and function as incretin mimetics by promoting pancreatic insulin release (3). It may be possible to further increase the ability of anti-obesity drugs to improve metabolic health through combination with agents that target distinct molecular mechanisms that impinge on adipose tissue (AT) to promote energy expenditure and ameliorate fat dysfunction, hallmarks of which are insulin resistance, inflammation and fibrosis (4). In this regard, there is interest in developing drugs that directly augment brown AT (BAT) function and/or stimulate white AT (WAT) ‘beiging’ as a means of treating metabolic disease (5).

While the primary role of WAT is to store energy in the form of triglycerides in unilocular white adipocytes, brown adipocytes within BAT harbor small, multilocular lipid droplets and an abundance of mitochondria, which dissipate chemical energy as heat through non-shivering thermogenesis. Heat production by BAT is governed by uncoupling protein-1 (UCP1), which catalyzes mitochondrial proton leak and thereby uncouples electron transport from ATP synthesis (6-8). Beige adipocytes within WAT are similar to brown adipocytes in that they express UCP1, have multiple lipid droplets, and are mitochondria-rich (9). The existence of functional BAT in adult humans was confirmed by a series of positron-emission tomography (PET) studies, and body mass index and percent body fat were found to negatively correlate with BAT abundance, suggesting the possibility that stimulating brown and beige adipocytes would be useful for the treatment of obesity and obesity-related disorders. (10-13).

Pharmacological approaches to stimulate BAT and beiging of WAT in humans have focused on the use β_3_-adrenergic receptor (β_3_-AR) agonists. β_3_-AR stimulation directly enhances lipolysis and energy expenditure, and also triggers downstream signaling events that leads to potent induction of UCP1 expression in brown and white adipocytes (14). However, while β_3_-AR agonists were shown to acutely increase energy expenditure and insulin sensitivity in humans, they have persistently failed to promote weight loss upon chronic administration (15-18). More recent human studies have employed mirabegron, a β_3_-AR agonist that is approved by the US Food and Drug Administration (FDA) for the treatment of overactive bladder. Acute administration of mirabegron was shown to stimulate BAT metabolic activity in humans, but typically required high doses of the agonist that resulted in cardiovascular side effects such as tachycardia and hypertension (19-22). In healthy young adults or older, obese and insulin resistant humans, chronic mirabegron treatment improved glucose tolerance, but the drug only increased BAT activity in healthy subjects (23-25). The variable and limited effects of mirabegron on BAT activity in humans could be due to a greater reliance on β_1_-AR and β_2_-AR signaling in human versus murine AT, where the β_3_-AR predominates (20,26). Alternatively, the restricted efficacy of mirabegron could be a consequence of ‘catecholamine resistance’, wherein chronic exposure to ligand and/or inflammatory cues results in a reduction in β_3_-AR expression and signaling (27).

Histone deacetylase 11 (HDAC11) is a zinc-dependent enzyme that, paradoxically, functions as a robust lysine demyristoylase, with a catalytic efficiency >10,000-fold higher for myristoyl-lysine versus acetyl-lysine (28-30). Whole-body deletion of HDAC11 in mice prevented weight gain and improved overall metabolic health in the face of chronic high-fat feeding, and the beneficial effects of HDAC11 knockout were linked to increased UCP1 expression in BAT and WAT (31,32). Subsequent work revealed a cytoplasmic function for HDAC11 in the control of adipocyte β_3_-AR signaling. By demyristoylating two lysine residues within the protein kinase A (PKA) binding domain of the scaffolding protein, A kinase–anchoring protein 12 (AKAP12)/gravin-α, HDAC11 prevents gravin-α:β-AR:PKA complexes from localizing to membrane lipid rafts that are required to stimulate downstream signaling that culminates in UCP1 expression (33). However, whether knockout and/or pharmacological inhibition of HDAC11 in adipocytes is sufficient to drive UCP1 expression independently of β-ARs remained known.

Here, we demonstrate that adipocyte-specific knockout of HDAC11 in mice profoundly induces expression of UCP1 in BAT and WAT, resulting in elevated body temperature. These effects are recapitulated by treating mice *in vivo* or human AT *ex vivo* with a pharmacological inhibitor of HDAC11. We provide evidence for biphasic, gravin-α myristoylation-dependent induction of UCP1 protein expression, with an early post-transcriptional response that is independent of PKA, and a later response requiring PKA and *de novo Ucp1* mRNA synthesis. HDAC11 inhibition stimulates UCP1 expression even in the setting of catecholamine resistance where β-AR signaling is blocked. These findings define cell autonomous, multi-modal roles for HDAC11 in the repression of UCP1 expression. Furthermore, the data underscore the potential of inhibiting HDAC11 to bypass catecholamine resistance and directly modulate AT phenotype in the context of obesity.

## Results

### Adipocyte-specific knockout of HDAC11 in mice is sufficient to induce UCP1 protein expression and promote thermogenesis

Global deletion of HDAC11 in mice enhances UCP1 expression in AT (31,32). To address whether HDAC11 is a cell autonomous repressor of thermogenic gene expression, mice harboring a conditional allele with *loxP* sites flanking exon 2 of the *Hdac11* gene were generated (Figure 1A). *Hdac11^fl/fl^* mice were crossed with mice containing a transgene of the adiponectin promoter driving constitutive expression of Cre recombinase (*Adipoq-Cre*) to yield adipocyte-specific *Hdac11* conditional knockout mice (*Hd11^cKO^*) (Figure 1, B - D). HDAC11 protein levels were greatly reduced in the *Hd11^cKO^* epididymal WAT (eWAT), inguinal WAT (ingWAT) and BAT compared to wildtype (WT) controls (Figure 1, E and F). Genetic deletion of HDAC11 in adipocytes led to marked induction of UCP1 protein expression in each of these AT depots (Figure 1, E - G).

**Figure 1.**
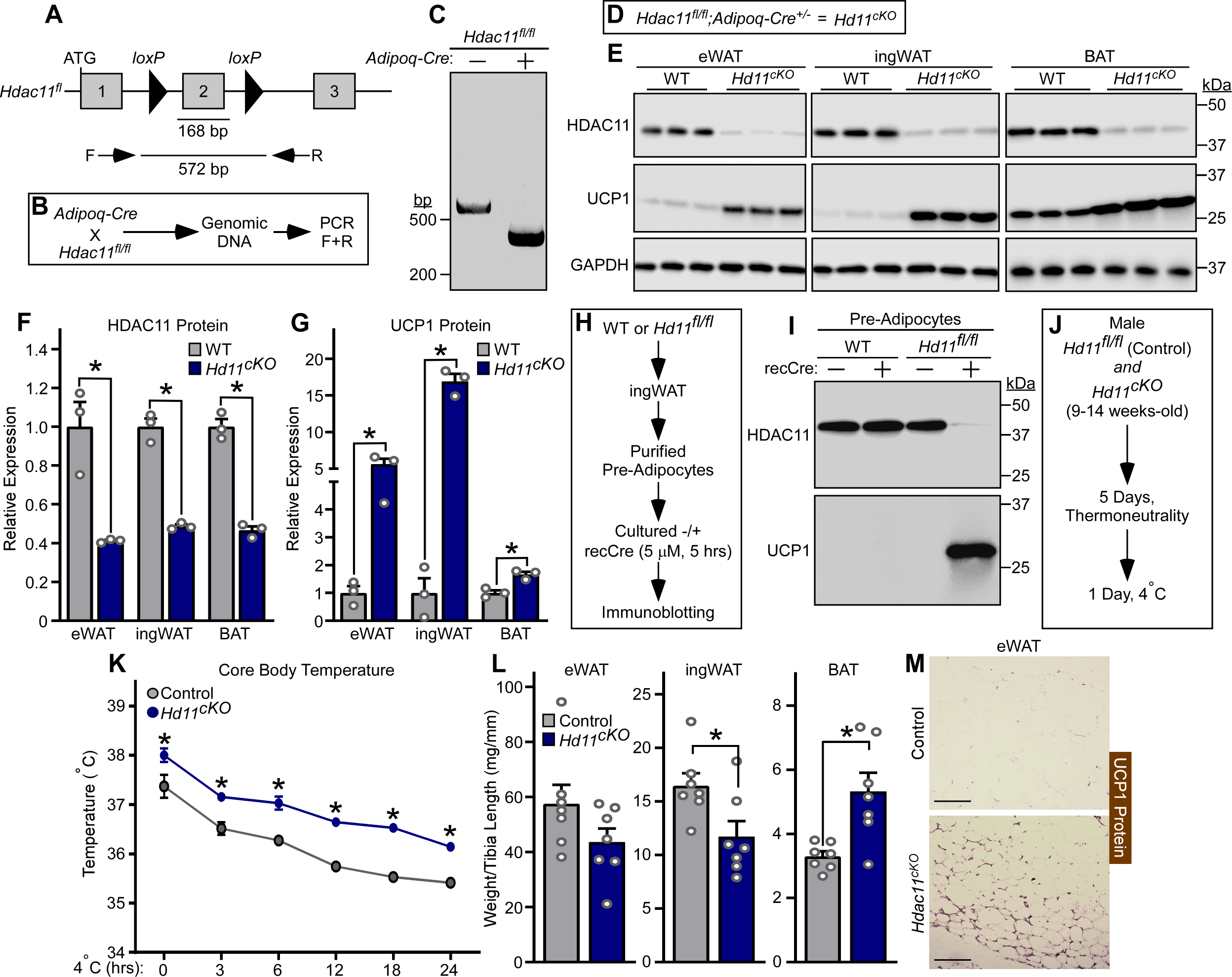
Adipocyte-specific knockout of HDAC11 in mice is sufficient to induce UCP1 protein expression and promote thermogenesis. **(A)** Schematic representation of the *Hdac11* floxed allele (HDAC11^fl^), with *loxP* sites flanking the 162 base pair (bp) exon 2; the predicted size of the resulting PCR product generated from genomic DNA and forward (F) and reverse (R) primers is shown. **(B)** Schematic representation of the cross between transgenic mice harboring an adiponectin promoter-driven Cre recombinase (*AdipoQ-Cre*) and *Hdac11^fl/fl^* mice and the genomic DNA PCR approach to assessing excision of *Hdac11* exon 2. **(C)** Agarose gel image of genomic DNA PCR products from *Hdac11^fl/fl^* mice without (-) and with (+) the *AdipoQ-Cre* transgene. (**D**) Diagram describing the genotype and definition of the adipocyte-specific conditional *Hdac11* cKO mouse (*Hd11^cKO^*). (**E**) HDAC11 and UCP1 protein levels were assessed by immunoblotting with homogenates of epididymal white adipose tissue (eWAT), inguinal white adipose tissue (ingWAT) and interscapular brown adipose tissue (BAT) from 10-12-week-old wildtype (WT) and *Hdac11^cKO^* mice. GAPDH served as a loading control; n=3 biological replicates/group. (**F** and **G**) Densitometric analysis of HDAC11 and UCP1 protein expression in (**E**), normalized to GAPDH and plotted relative to WT controls. Data are presented as mean +SEM, with \**P*<0.05 as determined by 2-tailed, unpaired *t* test. (**H**) Schematic representation of the cell culture experiment with primary inguinal pre-adipocytes isolated from WT or *Hdac11*^fl/fl^ mice and treated with recombinant, cell-permeable Cre recombinase. (**I**) Immunoblot analysis of HDAC11 and UCP1 protein expression in pre-adipocytes. (**J**) Schematic representation of the 24-hour 4°C challenge experiment. (**K**) Core body temperature of was determined at the indicated times. Data are depicted as mean ±SEM, with \**P*<0.05 vs. WT mice at a given time as determined by two-way ANOVA with Sidak’s multiple comparisons test; n=7 biological replicates/group. (**L**) Adipose tissue weight, normalized to tibia length, determined upon necropsy after the 24-hour 4°C challenge. Data are depicted as mean +SEM, with \**P*<0.05 as determined by 2-tailed, unpaired *t* test. (**M**) Immunohistochemistry images of UCP1 protein expression in eWAT from mice sacrificed following the 24-hour 4°C challenge; scale bar = 200 μm.

To further address cell autonomous functions of HDAC11, preadipocytes were isolated from the stromal vascular fraction of ingWAT from WT and *Hdac11^fl/fl^* mice and cultured in the absence or presence of recombinant cell-permeable Cre recombinase (Figure 1H). Remarkably, incubation of *Hd11^fl/fl^* preadipocytes with Cre for only 5 hours led to a dramatic reduction in HDAC11 protein abundance, which correlated with strong induction of UCP1 protein expression (Figure 1I).

As UCP1 functions to facilitate non-shivering thermogenesis, differences in core body temperature were examined in *Hd11^c^*^KO^ and *Hd11^fl/fl^* control mice exposed to 4 °C for 24 hours following five days of acclimatization at thermoneutrality (28 - 30 °C) (Figure 1J). Compared to controls, *Hd11^c^*^KO^ mice had higher core body temperature at baseline and throughout the 24-hour cold challenge (Figure 1K). Analysis of AT following necropsy revealed decreased WAT and increased BAT abundance and evidence of WAT beiging in *Hd11^cKO^* mice compared to controls (Figure 1, L and M). Together, these findings establish that HDAC11 serves an adipocyte autonomous role in the control of AT thermogenic protein expression and remodeling.

### Selective pharmacological inhibition of HDAC11 in mice is sufficient to induce thermogenesis and UCP1 expression

To address the therapeutic potential of inhibiting HDAC11 to alter AT phenotype, mice were treated with a highly selective inhibitor of HDAC11, FT895, or vehicle control, for five days at thermoneutrality prior to a 24-hour 4 °C challenge (Figure 2A). Mice pretreated with FT895 had a higher core body temperature at baseline and throughout the 4 °C challenge (Figure 2B). Furthermore, compared to controls, mice treated with FT895 had reduced WAT and increased BAT abundance, and had profoundly augmented expression of UCP1 protein in eWAT, ingWAT and BAT (Figure 2, C - E). These findings illustrate that pharmacological inhibition of HDAC11 catalytic activity phenocopies genetic deletion of HDAC11 in adipocytes, resulting in enhanced thermogenesis in association with strong induction of UCP1 protein expression and reduced remodeling of both WAT and BAT.

**Figure 2.**
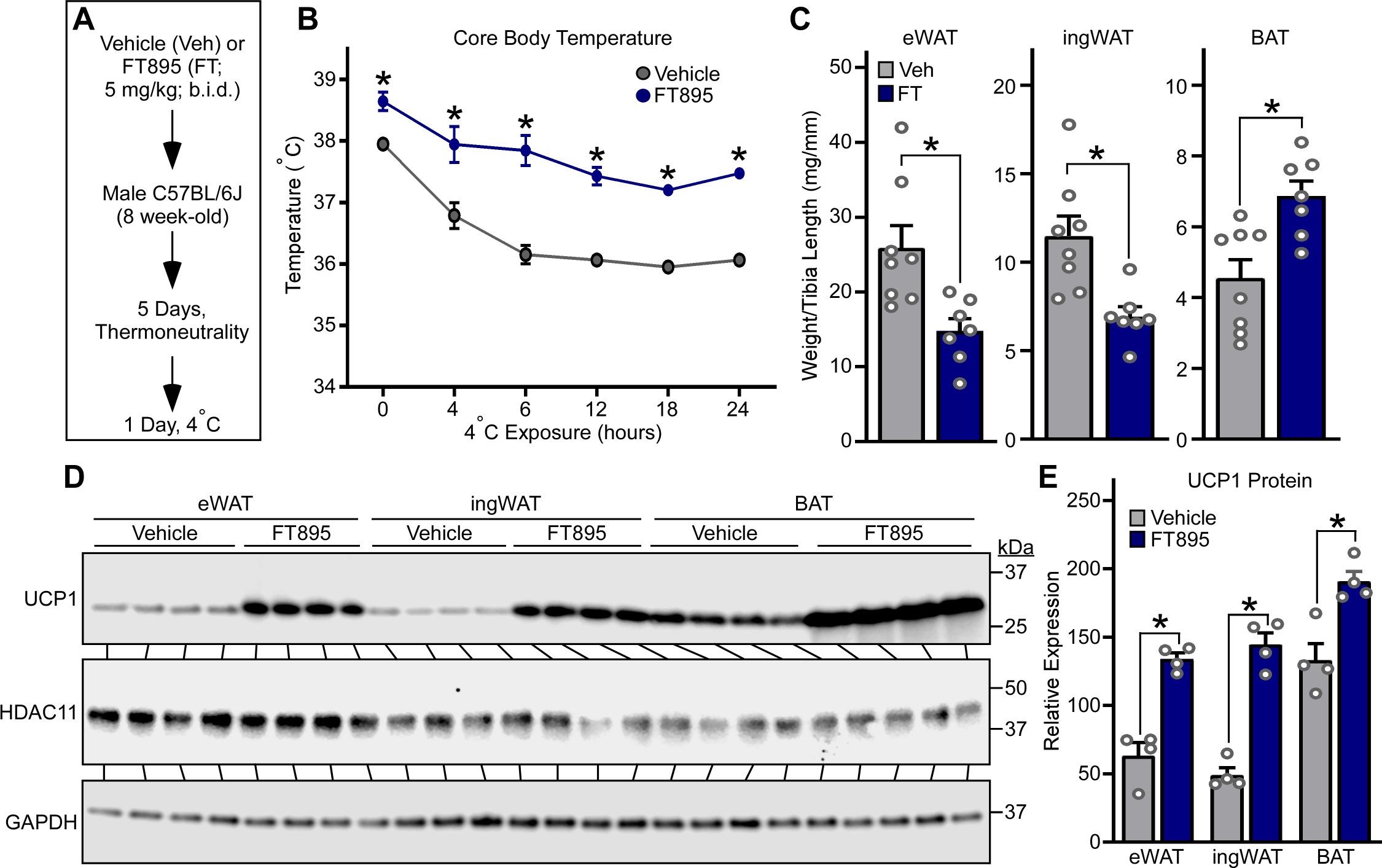
Selective pharmacological inhibition of HDAC11 in mice is sufficient to induce thermogenesis and UCP1 expression. **(A)** Schematic representation of the 24-hour 4°C challenge experiment employing the selective HDAC11 inhibitor, FT895. **(B)** Core body temperature was determined at the indicated times. Data are depicted as mean ±SEM, with \**P* < 0.05 vs. vehicle-treated mice at a given time as determined by two-way ANOVA with Sidak’s multiple comparisons test; vehicle (n=8 biological replicates) and FT895 (n=7 biological replicates). (**C**) Epididymal white adipose tissue (eWAT), inguinal white adipose tissue (ingWAT) and interscapular brown adipose tissue (BAT) weights, normalized to tibia length, determined upon necropsy after the 24-hour 4°C challenge. Data are depicted as mean +SEM, with \**P*<0.05 as determined by 2-tailed, unpaired *t* test. (**D**) HDAC11 and UCP1 protein levels were assessed by immunoblotting with homogenates of AT obtained from mice sacrificed after the 24-hour 4°C challenge; n=4 biological replicates/group. (**E**) Densitometric analysis of HDAC11 and UCP1 protein in (**D**), normalized to GAPDH. Data are depicted a mean +SEM, with \**P*<0.05 as determined by 2-tailed, unpaired *t* test.

### HDAC11 triggers biphasic induction of UCP1 protein expression through post-transcriptional and transcriptional mechanisms

Cultured adipocytes were used to elucidate the mechanism(s) by which HDAC11 inhibition induces UCP1 protein expression. Consistent with findings made with AT *in vivo* and *ex vivo*, treatment of differentiated murine white-like adipocytes, 3T3-L1 cells, with FT895 for 1 hour led to robust induction of UCP1 protein expression (Figure 3, A - C). Evaluation of primary adipocytes from WT and *Hdac11* knockout mice established that FT895 stimulates UCP1 expression by inhibiting HDAC11 as opposed to through an off-target action since, compared to WT cells, UCP1 abundance was elevated at baseline in *Hdac11* knockout adipocytes and not further increased by the inhibitor (Supplemental Figure 1). FT895 also stimulated UCP1 protein expression in HIB1B adipocytes, which exhibited higher basal UCP1 expression due to their brown adipocyte-like phenotype, and promoted expression of UCP1 in cultured primary human subcutaneous adipocytes (Figure 3, D - F).

**Figure 3.**
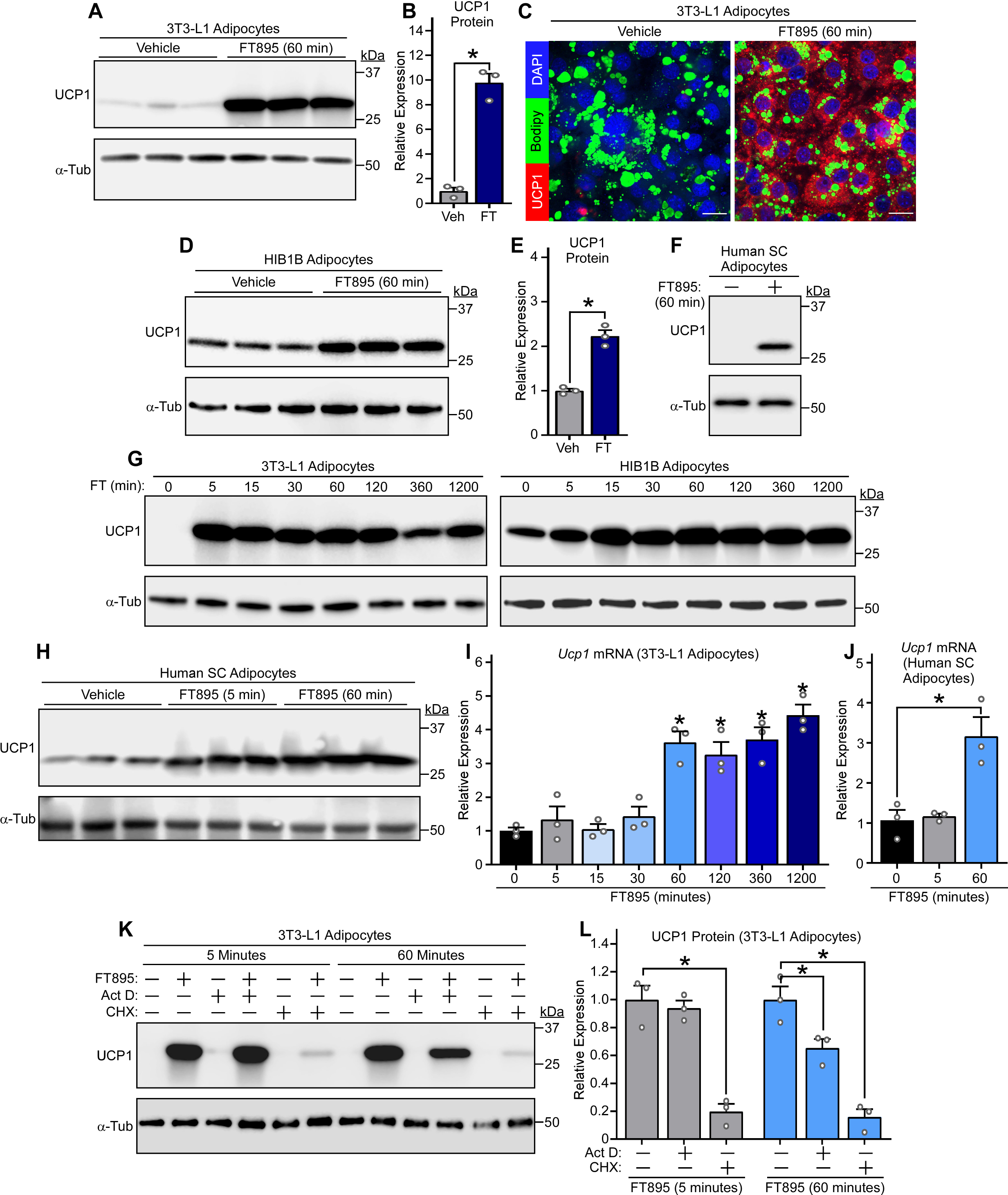
HDAC11 inhibition triggers biphasic induction of UCP1 protein expression through post-transcriptional and transcriptional mechanisms. **(A)** Immunoblot analysis of UCP1 protein in 3T3-L1 white adipocytes treated with vehicle control or FT895 for 60 minutes; α-tubulin (α-Tub) served as a loading control; n=3 technical replicates/condition. **(B)** Densitometric analysis of UCP1 expression in **(A)**, normalized to α-tubulin and plotted as fold-change relative to untreated controls. Data are presented as mean +SEM, with \**P*<0.05 (2-tailed, unpaired *t* test). **(C)** Indirect immunofluorescence analysis of UCP1 protein (red) in 3T3-L1 adipocytes treated with vehicle control or FT895 for 60 minutes. Nuclei and lipid droplets were co-stained using DAPI (blue) and BODIPY (green), respectively; scale bar = 10 µm. **(D)** Immunoblot analysis with homogenates of HIB1B brown adipocytes treated with vehicle control or FT895 for 60 minutes; n=3 technical replicates/condition. **(E)** Densitometric analysis of UCP1 expression in **(D)**, normalized to α-tubulin and plotted as fold-change relative to untreated controls. Data are presented as mean +SEM, with \**P*≤0.05 (2-tailed, unpaired *t* test). **(F)** Immunoblot analysis with homogenates of cultured human subcutaneous (SC) adipocytes treated with FT895 for 60 minutes. **(G)** Immunoblot analysis with homogenates of 3T3-L1 adipocytes (left) and HIB1B adipocytes (right) treated with FT895 for the indicated times. **(H)** Immunoblot analysis with homogenates of cultured human SC adipocytes treated with FT895 for 5 and 60 minutes; n=3 technical replicates/condition. *Ucp1* mRNA expression in 3T3-L1 adipocytes (I) or human SC adipocytes (J) treated with FT895 for the indicated times was determined by qRT-PCR. Data were normalized to *18S* rRNA and are plotted as fold-change relative to the 0-minute point. Data are presented as mean +SEM, \**P*<0.05 vs. the 0-minute point as determined by one-way ANOVA with Tukey’s multiple comparisons test; n=3 technical replicates/condition. **(K)** 3T3-L1 adipocytes were pre-treated with vehicle (-), actinomycin D (Act D) or cycloheximide (CHX) for 30 minutes prior to exposure to FT895 for 5 or 60 minutes. Cells were homogenized and immunoblotting was performed. **(L)** Densitometric analysis of UCP1 expression in **(K)** as well as from two additional independent experiments (blots not shown), normalized to α-tubulin and plotted as fold-change relative to FT895 treatment alone. Data are presented as mean +SEM, with \**P*<0.05 vs. FT895 treatment as determined by one-way ANOVA with Tukey’s multiple comparisons test.

Time course experiments were performed to define the kinetics of UCP1 induction following HDAC11 inhibition. UCP1 protein expression was dramatically upregulated as early as 5 minutes following FT895 treatment and was sustained 20 hours-post-exposure to the HDAC11 inhibitor in both 3T3-L1 and HIB1B adipocytes (Figure 3G). Rapid induction of UCP1 protein expression was also observed in primary human subcutaneous adipocytes exposed to FT895 (Figure 3H). Parallel gene expression analyses with 3T3-L1 and human adipocyte homogenates revealed that FT895 stimulated *Ucp1* mRNA expression beginning 60 minutes following treatment, suggesting that UCP1 protein induction following 5 minutes of FT895 exposure occurs through a post-transcriptional mechanism (Figure 3, I and J). To address this possibility, studies were performed with actinomycin D, which blocks gene transcription, and cycloheximide, which suppresses protein translation. Pretreatment with actinomycin D had no effect on UCP1 protein induction following 5 minutes of FT895 treatment, but significantly reduced UCP1 protein abundance following 60 minutes of HDAC11 inhibition (Figure 3, K and L). In contrast, cycloheximide pretreatment strongly reduced UCP1 induction at both time points (Figure 3, K and L). These data suggest that HDAC11 inhibition stimulates adipocyte UCP1 protein expression in a biphasic and bimodal manner, with acute induction occurring through a post-transcriptional mechanism, and delayed induction requiring *de novo Ucp1* mRNA synthesis.

### Biphasic UCP1 induction upon HDAC11 inhibition is dependent on gravin-α lysine myristoylation

Myristoylation of the anchoring protein, gravin-α, on lysine residues 1502 and 1505 drives gravin-α:β_3_-AR complexes into caveolin-rich lipid rafts to promote downstream PKA signaling (33). Conversely, demyristoylation of gravin-α by HDAC11 blocks β_3_-AR signaling (Figure 4A). A click chemistry method with biotinylated myristic acid alkyne (Alk-12) confirmed that gravin-α was myristoylated in 3T3-L1 adipocytes following treatment with FT895 for 5 and 60 minutes (Figure 4, B and C).

**Figure 4.**
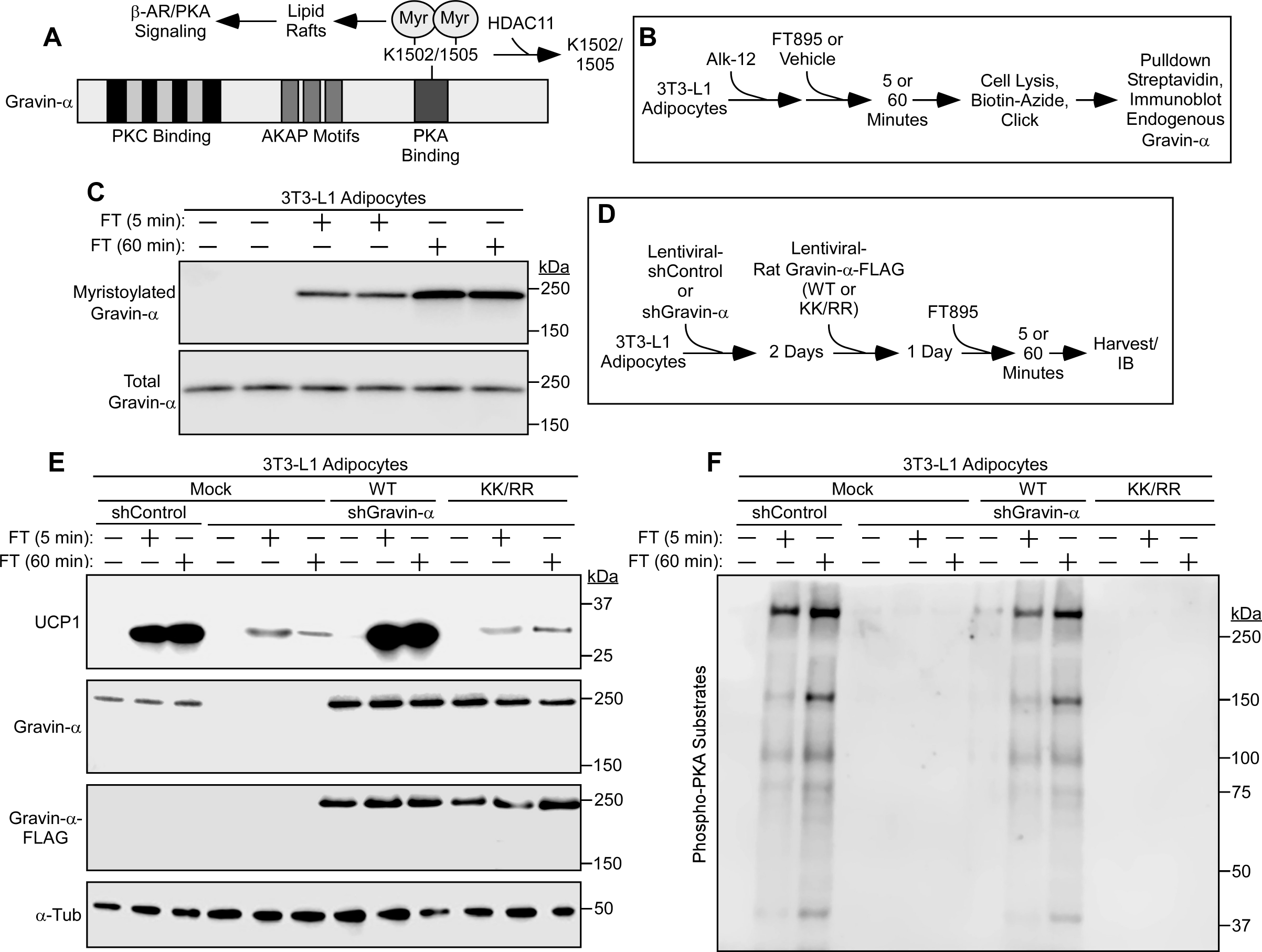
Biphasic UCP1 induction upon HDAC11 inhibition is dependent on gravin−α lysine myristoylation. (**A**) Schematic representation of gravin-α protein structure, indicating the two lysine residues that are demyristoylated by HDAC11. Myristoylation of these two conserved lysines upon HDAC11 inhibition drives gravin-α:β-adrenergic receptor (β-AR) complexes into membrane lipid rafts, resulting in downstream protein kinase A (PKA) signaling. (**B**) Schematic depiction of the click chemistry experiment employing a myristic acid ‘click’ tag (Alk-12) to determine if acute HDAC11 inhibition with FT895 promotes gravin-α myristoylation. (**C**) Immunoblot analysis to detect endogenous myristoylated gravin-α and total gravin-α. (**D**) Schematic depiction of the experiment to determine if gravin-α and its myristoylation are required for induction of UCP1 protein expression following HDAC11 inhibition with FT895. (**E**) Immunoblot analysis to detect UCP1, total gravin-α and FLAG-tagged gravin-α protein expression; α-tubulin (α-Tub) served as a loading control. (**F**) Immunoblot analysis with an antibody that recognizes proteins containing phospho-serine/threonine residues within a consensus PKA target site (RRXS*/T*).

To address the role of gravin-α and its myristoylation in the control of acute and chronic UCP1 protein induction following HDAC11 inhibition, endogenous gravin-α expression was knocked down in mouse 3T3-L1 adipocytes using short hairpin RNA, and WT or a myristoylation-resistant version of rat gravin-α (K1502/1505R; [KK/RR]) were subsequently ectopically expressed in the cells (Figure 4D). Knockdown of endogenous gravin-α dramatically reduced UCP1 protein induction following 5 and 60 minutes of FT895 treatment, and addback of WT, but not KK/RR, rescued UCP1 expression (Figure 4E). These findings correlated with stimulation of PKA signaling in the cells, with 5 and 60 minutes of FT895 treatment promoting PKA substrates phosphorylation in a manner that was dependent on site-specific myristoylation of gravin-α (Figure 4F).

### Inhibition of HDAC11 stimulates PKA-independent and PKA-dependent induction of UCP1 expression

To determine if acute and delayed UCP1 protein induction following HDAC11 inhibition are dependent on PKA signaling, which is the canonical pathway for stimulation of thermogenic gene expression, 3T3-L1 cells were treated with FT895 in the absence or the presence of the PKA inhibitor, H89 (Figure 5A). H89 administration had a limited effect on UCP1 protein induction following 5 minutes of FT895 treatment, but dramatically suppressed UCP1 expression in cells treated with the HDAC11 inhibitor for 60 minutes (Figure 5, B and C). Immunoblotting with an anti-phospho PKA substrates antibody confirmed the ability of H89 to effectively block FT895-mediated induction of general PKA signaling at both time points (Figure 5D). Analysis of RNA from independent samples revealed that H89 blocked FT895-mediated induction of *Ucp1* mRNA expression following 60 minutes of treatment; consistent with prior results, no induction of *Ucp1* mRNA expression was noted after 5 minutes of treatment with the HDAC11 inhibitor (Figure 5E). Thus, although induction of UCP1 protein expression following treatment with FT895 for 5 and 60 minutes is dependent on gravin-α myristoylation, and PKA is activated by the HDAC11 inhibitor at each time point, only the 60-minute response requires the activity of the kinase. This requirement is likely due to PKA-mediated induction of *Ucp1* mRNA synthesis.

**Figure 5.**
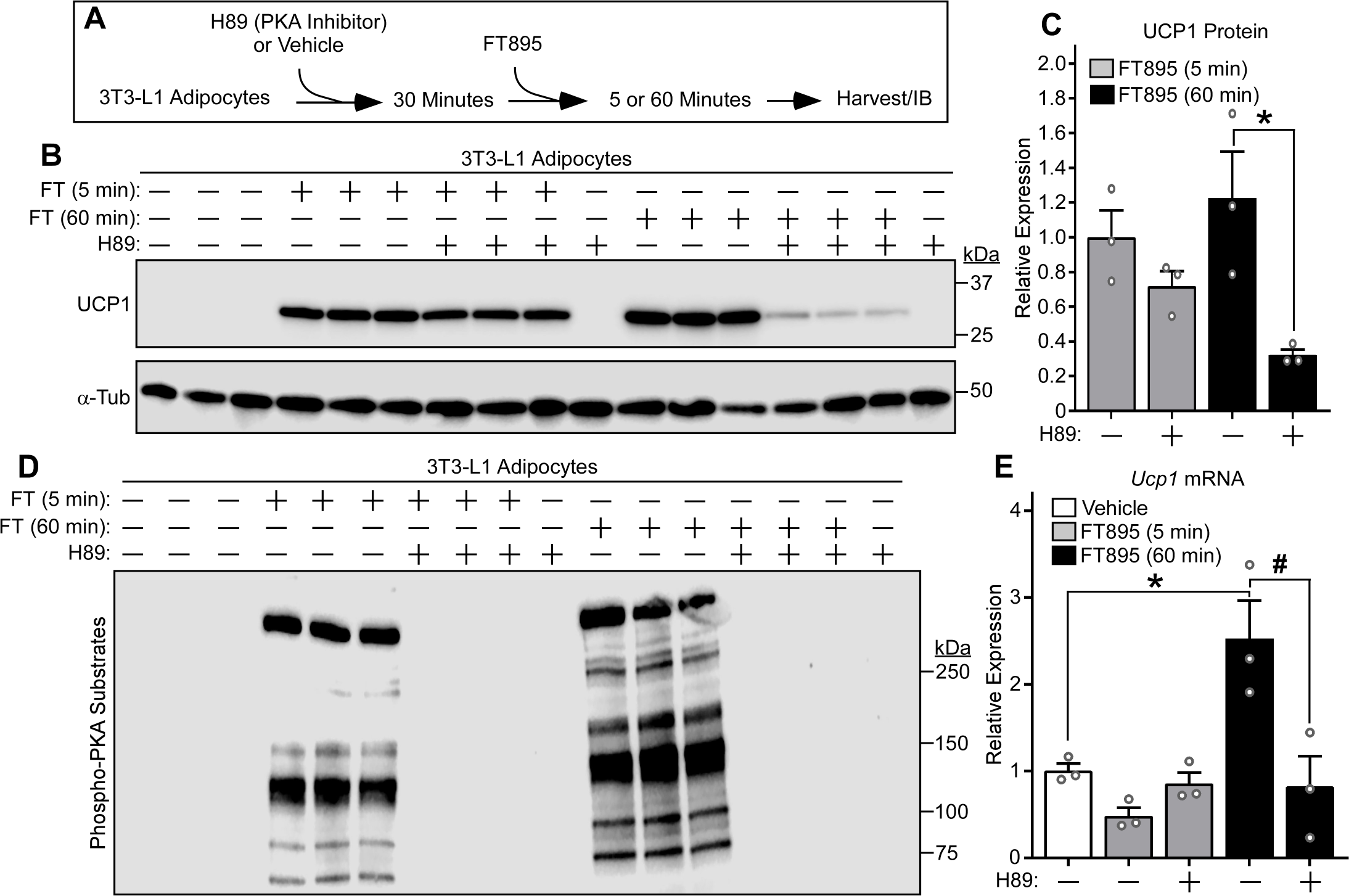
Inhibition of HDAC11 stimulates PKA-independent and PKA-dependent induction of UCP1 expression. (**A**) Schematic representation of the experiment to test the dependency of FT895-induced UCP1 expression on PKA signaling. (**B**) Immunoblot analysis of UCP1 protein expression, with α-tubulin (α-Tub) as a loading control; n=3 technical replicates/condition. (**C**) Densitometric analysis of the UCP1 signal in (**B**), normalized to α-Tub and depicted as fold-change relative to cells treated with FT895 alone. Data are presented as mean +SEM, with \**P* < 0.05 as determined by 2-tailed, unpaired *t* test. (**D**) 3T3-L1 homogenates were immunoblotted with an antibody that recognizes proteins containing phospho-serine/threonine residues within a consensus PKA target site (RRXS*/T*). (**E**) *Ucp1* mRNA expression in 3T3-L1 adipocytes treated with FT895 for the indicated times in the absence or presence of H89 pretreatment was determined by qRT-PCR. Data were normalized to *18S* rRNA and are plotted as fold-change relative to the vehicle treated cells. Data are presented as mean +SEM, \**P* < 0.05 vs. vehicle treatment and ^#^*P*<0.05 vs 60-minute FT895 treatment without H89 as determined by one-way ANOVA with Tukey’s multiple comparisons test; n=3 technical replicates/condition.

### HDAC11 inhibition triggers UCP1 induction and PKA signaling in murine and human models of adipocyte catecholamine resistance

Although the β_3_-AR is not regulated by classical phosphorylation-dependent receptor desensitization, recent work has demonstrated that expression of the receptor is downregulated in response to chronic ligand exposure, nutrient excess and/or inflammatory cues, leading to AT catecholamine resistance (27). Given the ability of HDAC11 inhibition to elicit UCP1 expression and PKA signaling independently of β-AR receptor ligand binding, experiments were performed to address the possibility that FT895 treatment circumvents β_3_-AR downregulation in cell-based and *in vivo* models of catecholamine resistance. Initially, 3T3-L1 adipocyte were exposed to DMSO vehicle or the β_3_-AR agonist CL-316,243 for 20 hours prior to 1 hour of re-exposure to these agents, forskolin, which served as a positive control by virtue of its ability to stimulate adenylyl cyclase downstream of the receptor, or FT895 (Figure 6A). Consistent with prior findings, chronically treating adipocytes with CL-316,243 led to reduced β_3_-AR protein expression and blocked subsequent UCP1 induction and PKA activation in response to acute re-exposure to this agonist (Figure 6, B and C) (27). Strikingly, FT895 was as effective as forskolin at bypassing β_3_-AR downregulation to stimulate UCP1 protein expression in adipocytes chronically treated with CL-316,243, and also promoted PKA signaling in these cells (Figure 6, B and C).

**Figure 6.**
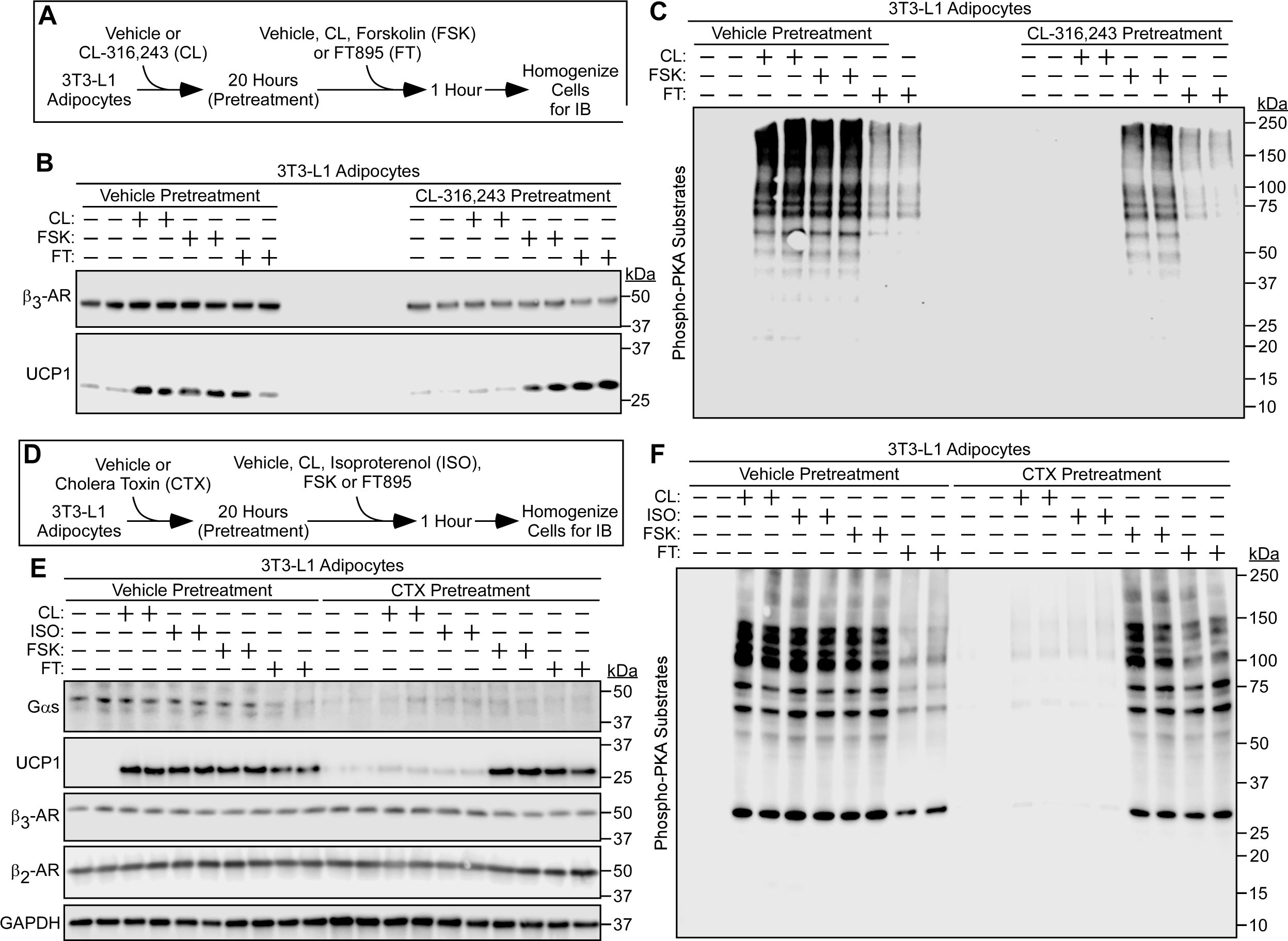
HDAC11 inhibition promotes UCP1 expression and PKA signaling in cell-based models of catecholamine resistance. (**A**) Schematic representation of the cell culture experiment to determine if HDAC11 inhibition promotes adipocyte UCP1 expression and PKA signaling in the context of downregulated β_3_-adrenergic receptor (β_3_-AR) expression due to chronic agonist exposure. (**B**) Immunoblot analysis with homogenates of 3T3-L1 cells pretreated with vehicle or the β_3_-AR agonist CL-316,243 (CL) for 20 hours followed by 1-hour treatment with vehicle (-), CL, the adenylyl cyclase activator forskolin (FSK) or FT895; n=2 technical replicates/condition. (**C**) These same samples were independently immunoblotted with an antibody that recognizes proteins containing phospho-serine/threonine residues within a consensus PKA target site (RRXS*/T*). (**D**) Schematic representation of the cell culture experiment to determine if HDAC11 inhibition promotes adipocyte UCP1 expression and PKA signaling in the context of catecholamine resistance due to chronic cholera toxin (CTX) exposure. (**E**) Immunoblot analysis of the indicated proteins in homogenates of 3T3-L1 cells pretreated with vehicle or the β_3_-AR agonist CL-316,243 (CL) for 20 hours followed by 1-hour treatment with vehicle (-), the non-selective β-AR agonist isoproterenol (ISO), CL, the adenylyl cyclase activator forskolin (FSK) or FT895; n=2 technical replicates/condition. (**F**) These same samples were independently immunoblotted with the anti-phospho-PKA substrates antibody.

Signals emanating from the β_3_-AR are transduced by the heterotrimeric G protein, G_s_ (14). A complementary cell-based model of adipocyte catecholamine resistance took advantage of the fact that, while acute treatment of cells with cholera toxin (CTX) stimulates adenylyl cyclase by catalyzing ADP-ribosylation of the alpha subunit of the heterotrimeric G protein, Gαs, chronic exposure to the toxin leads to downregulation of Gαs expression by promoting its degradation (Figure 6D) (34,35). Since the β_2_-AR also signals through Gαs, the CTX model enabled assessment of the possible contribution of this GPCR to the ability of FT895 to bypass CL-316,243 resistance seen above. Exposure of 3T3-L1 adipocytes to CTX for 12 hours led to reduced Gαs abundance without influencing β_3_-AR or β_2_-AR expression (Figure 6E). CTX pretreatment blocked subsequent UCP1 protein induction and PKA signaling following acute exposure of the cells to CL-316,243 or the non-selective β-AR agonist, isoproterenol. Conversely, forskolin and FT895 were still able to elicit these responses in cells treated with the toxin (Figure 6, E and F). Thus, HDAC11 inhibition is capable of promoting UCP1 protein expression and PKA activation independently of β-AR signaling in two independent cell-based models of catecholamine resistance, one in which β_3_-AR receptor expression is downregulated, and the other in which β_2_−and β_3_-AR coupling to Gαs is blocked.

To address the ability of HDAC11 inhibition to override AT catecholamine resistance *in vivo*, mice were injected with CL-316,243 or vehicle control for 12 hours prior to re-exposure to these agents or treatment with FT895 for an additional 1 hour (Figure 7A). Consistent with prior findings, CL-316,243 pretreatment led to a reduction in β_3_-AR protein expression in eWAT, ingWAT and BAT and blocked the subsequent induction of UCP1 protein and PKA signaling in each AT depot in response to acute re-exposure to the agonist (Figure 7, B - G) (27). Strikingly, FT895 was effective even in the setting of catecholamine resistance *in vivo*, promoting UCP1 protein expression and PKA signaling in eWAT, ingWAT and BAT of mice that were pretreated with CL-316-243 overnight (Figure 7, B - G).

**Figure 7.**
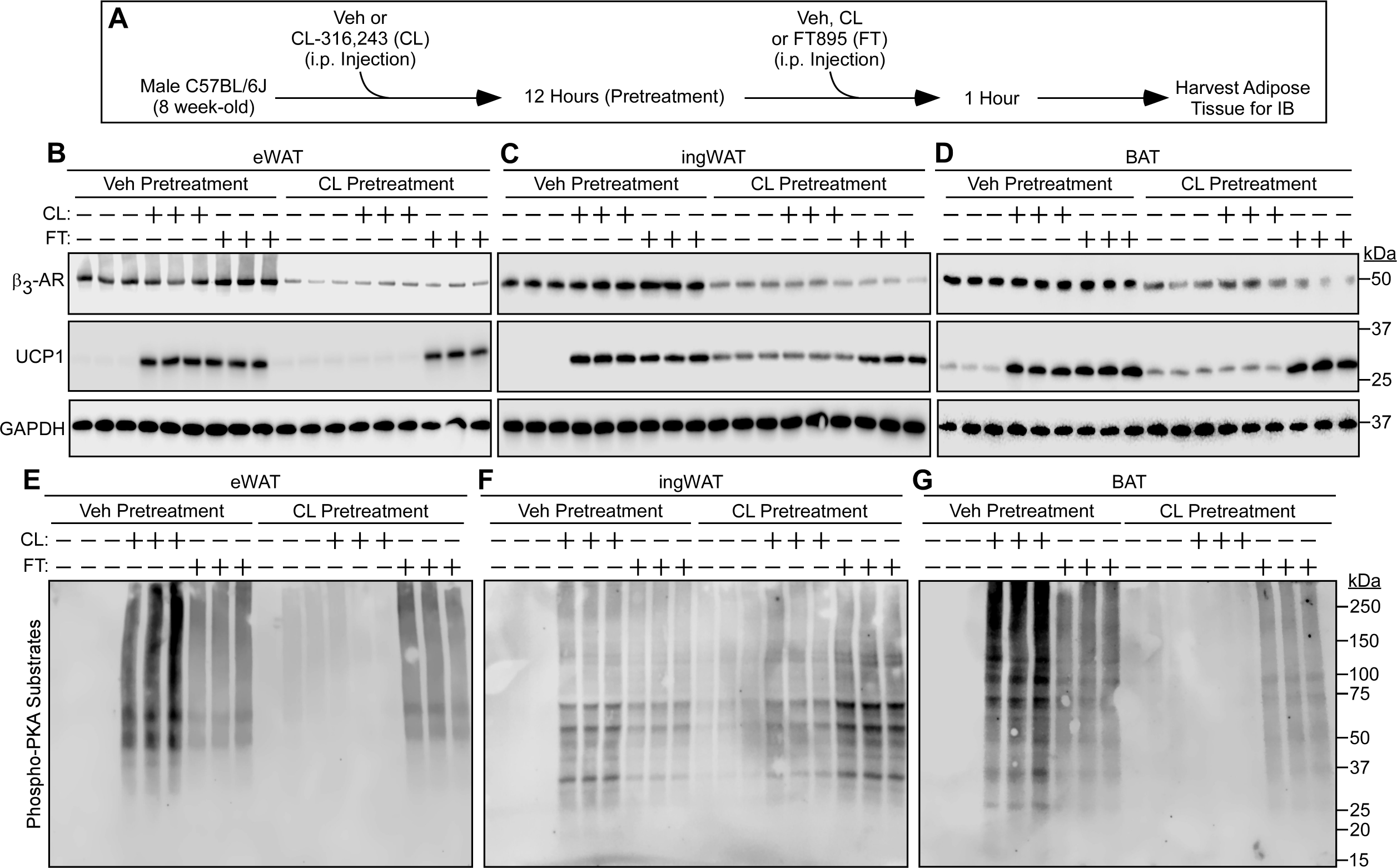
HDAC11 inhibition bypasses adipose tissue catecholamine resistance *in vivo*. (**A**) Schematic representation of the *in vivo* experiment to determine if HDAC11 inhibition promotes UCP1 expression and PKA signaling in the context of downregulated β_3_-adrenergic receptor (β_3_-AR) expression in adipose tissue. (**B** – **D**) Immunoblot analysis of the indicated proteins in homogenates of epididymal white adipose tissue (eWAT), inguinal white adipose tissue (ingWAT) and interscapular brown adipose tissue (BAT) in mice pretreated with vehicle (Veh) or CL-316,243 (CL) for 12 hours followed by 1-hour treatment with vehicle (-), CL or FT895 (FT); n=3 biological replicates/condition. (**E** – **G**) These same samples were independently immunoblotted with an antibody that recognizes proteins containing phospho-serine/threonine residues within a consensus PKA target site (RRXS*/T*).

### HDAC11 inhibition bypasses catecholamine resistance in human visceral adipose tissue ex vivo

To further address the translational potential of inhibiting HDAC11, human visceral adipose tissue (VAT) explants from obese individuals undergoing bariatric surgery were treated with FT895 or vehicle control *ex vivo*. BODIPY staining confirmed the integrity of the explants (Figure 8A). FT895 treatment of human VAT for just one hour was sufficient to induce UCP1 protein expression, which was maintained 20 hours following treatment (Figure 8B). Notably, FT895 stimulated UCP1 protein expression in human VAT as effectively the β_3_-AR agonist, CL-316,243, isoproterenol, or forskolin (Figure 8C).

**Figure 8.**
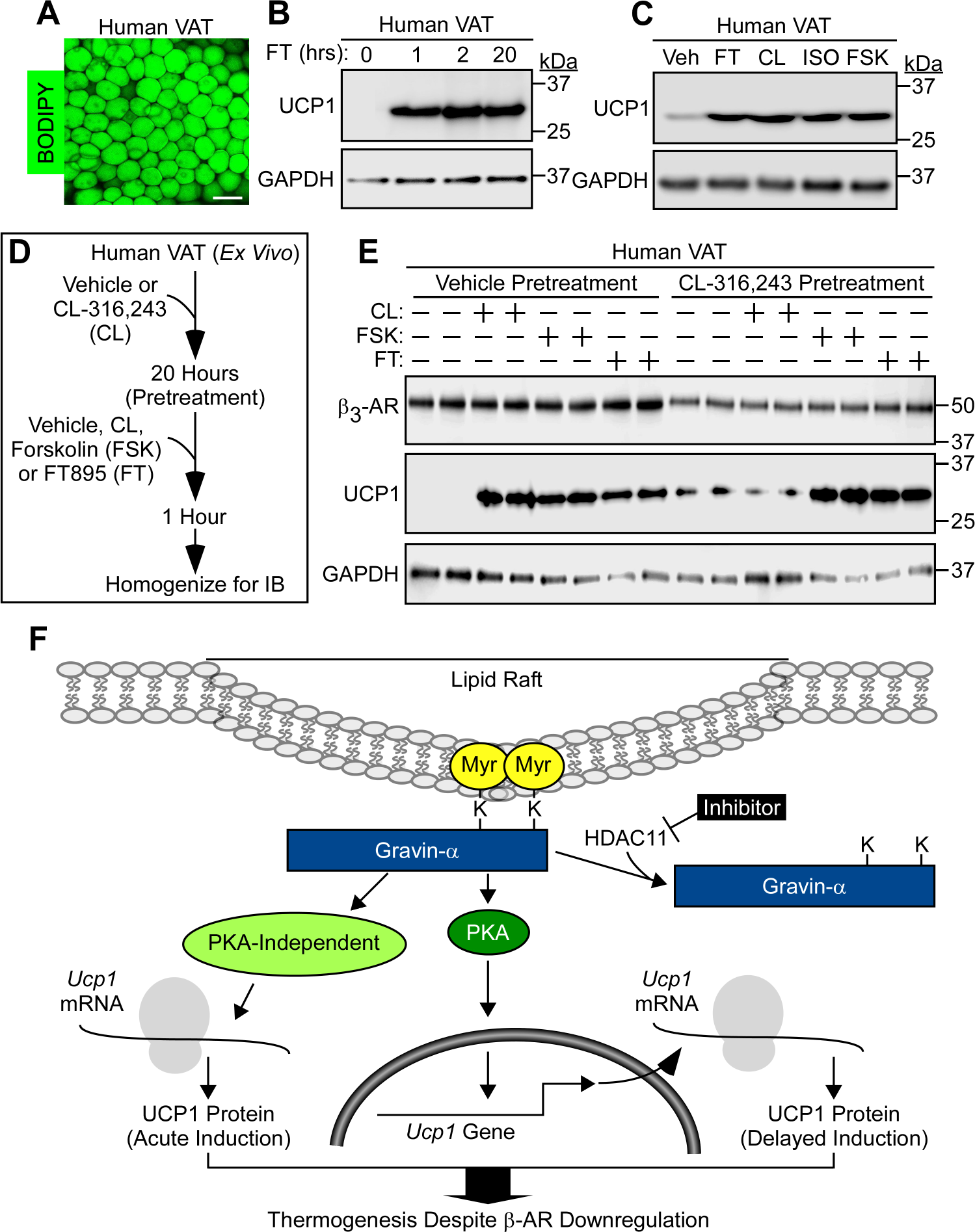
Therapeutic potential of inhibiting HDAC11 to circumvent adipocyte catecholamine resistance in humans. (**A**) Wholemount confocal microscopy image of human visceral adipose tissue (VAT) stained with BODIPY; scale bar = 100 µm. (**B**) Immunoblot analysis of UCP1 protein in human VAT treated *ex vivo* with the HDAC11 inhibitor, FT895, for the indicated times. GAPDH served as a loading control. **(C)** Immunoblot analysis of UCP1 and GAPDH in human VAT treated *ex vivo* with FT895 (FT; 1hr), the β_3_-AR agonist, CL-316,243 (CL; 3hrs), the non-selective β-AR agonist, isoproterenol (ISO; 2hrs), or the adenylyl cyclase activator, forskolin (FSK; 2hrs). **(D)** Schematic representation of the *ex vivo* experiment to determine if HDAC11 inhibition promotes UCP1 expression in the context of downregulated β_3_-adrenergic receptor (β_3_−AR) expression in human visceral adipose tissue (VAT). (**E**) Immunoblot analysis of the indicated proteins in homogenates of human VAT pretreated *ex vivo* with vehicle or the β_3_-AR agonist CL-316,243 (CL) for 20 hours followed by exposure to vehicle (-), CL, the adenylyl cyclase activator forskolin (FSK), or FT895 (FT) for 1 additional hour; n=2 biological replicates/condition. (**F**) A model for circumventing adipocyte catecholamine resistance through HDAC11 inhibition. HDAC11 normally demyristoylates two lysine residues in gravin-α, blocking the downstream signaling needed for UCP1 induction. When gravin-α is myristoylated on these lysines, such as upon HDAC11 inhibition with FT895, there is acute (5-minute) induction of UCP1 protein expression through a post-transcriptional mechanism that does not require PKA signaling, and there is delayed (60-minute) induction of UCP1 protein expression that requires new *Ucp1* mRNA synthesis and is dependent on PKA signaling. HDAC11 inhibition is capable of driving UCP1 protein expression even in the context of downregulated β-adrenergic receptors, suggesting the possibility of overcoming adipocyte catecholamine resistance in patients with metabolic disease through the use of selective HDAC11 inhibitors.

To assess whether HDAC11 inhibition is capable of promoting the thermogenic program in the setting of catecholamine resistance in human fat, VAT from two independent bariatric surgery patients was treated *ex vivo* with DMSO vehicle or CL-316,243 for 20 hours prior to re-exposure to these agents or treatment with forskolin or FT895 for an additional 1 hour (Figure 8D). Chronic treatment with the CL-316,243 led to reduced β_3_-AR protein expression and blocked subsequent UCP1 protein induction following acute agonist exposure (Figure 8E). Even in obese human AT, FT895 was comparable to forskolin at circumventing β_3_-AR downregulation to stimulate UCP1 protein expression, underscoring the therapeutic potential of inhibiting HDAC11 in the context of adipocyte catecholamine resistance (Figure 8E).

## Discussion

Attempts to stimulate BAT and beiging of WAT in humans using β_3_-AR agonists have met with only limited success, perhaps due to β_3_-AR downregulation in response to chronic ligand exposure and/or metabolic disease-induced inflammation, processes referred to as homologous and heterologous receptor desensitization, respectively (27). Here, we provide rationale for employing selective HDAC11 inhibitors to modulate AT phenotype independently of β_3_-AR agonism. Adipocyte-specific deletion or pharmacological inhibition of HDAC11 potently drives UCP1 expression and PKA signaling in BAT and WAT, even when the β_3_-AR is desensitized due to prolonged agonist treatment or when downstream signaling is blocked via CTX-mediated Gαs degradation. HDAC11 inhibition triggers ultra-rapid, post-transcriptional induction of UCP1 protein expression that occurs independently of PKA signaling, and also delayed induction of UCP1 expression that is governed by PKA and requires *de novo* mRNA synthesis (Figure 8F). We posit that this biphasic mechanism enables efficient thermogenic adaptation to acute and chronic environmental stress.

Stimulation of UCP1 expression following HDAC11 inhibitor treatment was dependent on site-specific myristoylation of gravin-α. Gravin-α is a scaffolding protein that harbors distinct binding domains for multiple proteins, including PKA and β_2_−and β_3_-ARs (33, 36-38). Previously, we showed that, upon HDAC11 inhibition, gravin-α is myristoylated on lysines 1502 and 1505, which leads to translocation of gravin-α:β_3_-AR:PKA protein complexes to plasma membrane lipid rafts containing proteins such as caveolin-1 and flotillin-2 (33). Targeting of the β_3_-AR to lipid rafts is required to efficiently trigger downstream PKA signaling and UCP1 induction in response to receptor ligands. The ability of FT895 to elicit these effects in the context of catecholamine resistance suggests that simply targeting gravin-α to lipid rafts is sufficient to promote downstream signaling, even in the absence of β_3_-AR engagement and Gs coupling. The mechanism(s) underlying such receptor-independent signaling mediated by myristoylated gravin-α remains unknown, but may involve recruitment of gravin-α-bound PKA to cyclic AMP rich regions juxtaposing the cell membrane, or lipid raft-dependent changes in association of gravin-α with effector proteins. We also cannot rule out the possibility that myristoylated gravin-α promotes adenylyl cyclase activity or alters phosphodiesterase function. In this regard, it is noteworthy that, at the level of phospho-PKA substrates immunoblotting, FT895 treatment elicited a pattern of protein phosphorylation that was qualitatively similar to that induced by forskolin or β-AR agonists.

The PKA-independent, gravin-α-dependent mechanism(s) for acute induction of UCP1 protein expression following FT895 treatment remains to be defined, but likely involves lipid raft-mediated activation of another gravin-α-associated signaling effector. While we are unaware of other demonstrations of a robust increase in UCP1 protein expression after 5 minutes of exposure to a stimulus, prior work illustrated a UCP1-dependent elevation of mouse BAT temperature within 2-3 minutes following administration of CL-316-243 *in vivo*, which is consistent with a role for post-transcriptional induction of UCP1 in acute adaptation to stress (39). Post-transcriptional mechanisms for regulating UCP1 protein expression independently of changes in *Ucp1* mRNA abundance have been described, and include translational control via binding of polyadenylation element binding protein 2 (CPEB2) and insulin-like growth factor 2 mRNA-binding protein 2 (IGF2BP2) to untranslated regions (UTRs) of the *Ucp1* transcript (40,41). Thus, it is possible that FT895-mediated activation of gravin-α signaling promotes acute UCP1 protein synthesis by modulating the activity of UTR binding proteins.

Transcriptional regulation of *Ucp1* gene has been studied in detail and involves binding of transcription factors such as cAMP response element binding protein (CREB), activating transcription factor 2 (ATF-2) and peroxisome proliferator-activated receptor gamma (PPARγ) to upstream distal enhancer and proximal promoter regions of the gene (42). PKA regulates *Ucp1* gene expression, in part, by simulating p38 mitogen-activated protein kinase, which phosphorylates ATF2 and PPARγ coactivator 1 alpha (PGC-1α), priming these proteins to bind to conserved cAMP and PPAR-associated response elements in the *Ucp1* enhancer region, respectively (43). Induction of adipocyte UCP1 protein expression following chronic treatment with FT895 presumably involves PKA-dependent activation of this signaling and transcription factor axis.

The *Ucp1* gene is also subject to active repression, such as through the action of nuclear receptor-interacting protein 1 (NRIP1) (44). Additionally, we previously demonstrated that a nuclear pool of HDAC11 interacts with the chromatin reader protein, BRD2, to suppress *Ucp1* gene expression (31). In this regard, it is noteworthy that FT895 failed to stimulate UCP1 expression in adipocytes in which gravin-α was knocked down, which was unexpected since the pharmacological inhibitor should engage BRD2-bound HDAC11 independently of the scaffolding protein. This finding suggests that inhibition of nuclear HDAC11 is necessary but not sufficient to induce UCP1 expression. We posit that in adipocytes chronically treated with FT895, gravin-α-dependent stimulation of PKA/p38 signaling provides a ‘second hit’ by activating transcription factors (e.g. ATF2 and PPARγ), which work in concert with inhibition of nuclear HDAC11 to promote *Ucp1* mRNA synthesis.

Surprisingly, deletion or pharmacological inhibition of HDAC11 in preadipocytes also led to UCP1 induction. Although uncommon, there are other examples of UCP1 expression in adipocyte precursor cells. Fibroblast growth factors 6 and 9 (FGF6/9) were found to function as adipokines with the capacity to stimulate UCP1 expression in preadipocytes (45). Upregulation of UCP1 by FGF6/9 occurred independently of canonical transcriptional mediators of thermogenesis such as PPARγ, and instead required a complex containing the nuclear hormone receptor, estrogen-related receptor-α. In a separate study, ectopic expression of the adipose tissue-enriched transcriptional co-activator, regulator PR domain containing 16 (PRDM16), in combination with forskolin treatment, was demonstrated to promote *Ucp1* mRNA expression in mouse embryonic fibroblasts (MEFs) (46). PRDM16-mediated expression of *Ucp1* expression in MEFs was in part dependent on stimulation of the transcriptional activity of another nuclear hormone receptor, thyroid hormone receptor. Whether HDAC11 inhibition activates either of the nuclear hormone receptor transcriptional pathways to promote UCP1 expression in preadipocytes remains to be determined.

Combining an HDAC11 inhibitor with existing anti-obesity drugs has the potential to enhance efficacy and improve the therapeutic index of these agents by directly altering AT phenotype. There are four FDA-approved HDAC inhibitors for the treatment of hematologic tumors, establishing the viability of targeting this class of enzymes to treat human diseases (47). Nonetheless, these ‘general’ HDAC inhibitors are poorly tolerated at the relatively high doses required to block tumor growth, and thus there is great interest in developing isoform-selective HDAC inhibitors with the goal of improving tolerability and efficacy. The discovery of FT895, which is >10,000-fold selective for HDAC11 over the other 10 zinc-dependent HDACs, has established the feasibility of selectively inhibiting this enzyme with a pharmacological agent (48). However, while FT895 is an excellent tool for research, it lacks oral bioavailability and harbors a hydroxamic acid warhead, which has been linked to mutagenicity in other compounds (48,49). Thus, future clinical development of HDAC11 inhibitors will require additional medicinal chemistry efforts to overcome these liabilities while maintaining selectivity and potency for this HDAC isoform.

In conclusion, we have shown that HDAC11 inhibition is as effective as β_3_-AR stimulation or direct adenylyl cyclase activation at stimulating UCP1 protein expression in murine and human AT. HDAC11 functions in an adipocyte-autonomous manner to control thermogenesis, as opposed to altering the phenotype of fat via indirect mechanisms such as by regulating adrenergic drive. HDAC11 inhibition triggers UCP1 expression through post-transcriptional and transcriptional mechanisms, both of which require site-specific myristoylation of gravin-α. The ability of HDAC11 inhibition to function in a β-AR-independent manner to bypass AT catecholamine resistance underscores the potential of targeting this lysine demyristoylase as a therapeutic strategy for metabolic diseases that have been recalcitrant to β_3_-AR agonists.

## Methods

### Further information can be found in Supplemental Methods

#### Creation of Hdac11 conditional knockout mice

CRISPOR and the Broad Institute sgRNA Design software were used to design guide RNA (50,51). Guides that performed best by both software programs were selected, and guide activity was verified by incubating guide RNA and Cas9 protein with a PCR product containing the target sequence and comparing the ratio of cut to uncut PCR product. Zygotes were injected with a single sgRNA (Synthego) with target sequence+PAM CAAGGCCCAAGGCCTGCACG + GGG, Cas9 protein (Integrated DNA Technologies, Alt-R® S.p. HiFi Cas9 Nuclease V3), and single stranded DNA template (Integrated DNA Technologies), starting with the weaker guide. Zygotes were then transferred into pseudopregnant recipients. F0 pups were genotyped by PCR using primers outside the region to be modified to identify putative positive founders; PCR primer sequences are found in Supplemental Table 1. These mice were then bred with appropriate WT mice. F1 male pups were genotyped and sequence confirmed for the desired introduction of a lone *loxP* site. F1 males were then used as sperm donors for IVF with B6N wildtype eggs and resulting zygotes were microinjected with second guide RNA with target sequence + PAM of CTCGGGGGACCTCCTATCTA + CGG, Cas9 protein, and a second DNA template to introduce the second *loxP* site. F0 mice were genotyped for both mutations. Mice with both *loxP* sites were bred with wildtype B6N mice. Co-segregation of the *loxP* sites indicates that they are in *cis*, indicative of a floxed allele. Once homozygous fl/fl mice were generated, both *loxP* sites were again sequenced via Sanger sequencing using primers outside of the DNA template sequence to verify the integration of fully intact *loxP* sites. *Hdac11^fl^* mice were crossed with *Adipoq-Cre* mice purchased from Jackson Labs (#028020). *Hdac11* floxed mice were generated in the Regional Mouse Genetics Core Facility, which is supported by both National Jewish Health and the University of Colorado Anschutz Medical Campus.

#### Cold challenge

All chemicals and reagents used for the current studies are described in Supplemental Table 2. Mice were housed singly for five days prior to commencement of the 24-hour 4 °C challenge. Core body temperature was monitored at the indicated times for the duration of the study using a rectal probe (Physitemp Instruments Inc.) connected to a physiological monitoring unit (THM150; VisualSonics). WT C57BL/6J mice (Jackson Labs, #000664) were employed for the 4°C studies with FT895. FT895 was delivered in a vehicle of 5% DMA/1% Tween 80/94% sterile water by i.p. injection for the five days of thermoneutral acclimatization and for the duration of the experiment. All 4°C challenge experiments were performed within the University of Colorado Nutrition and Obesity Center (NORC) small animal energy balance assessment core.

#### *In vivo model of* catecholamine *resistance*

The *in vivo* procedure to downregulate β_3_-AR expression in AT has been described previously (27). Briefly, 8-week old male WT mice (Jackson Labs, 000664) mice were pretreated with an i.p. injection of CL,316-243 (0.5 mg/kg in sterile saline) or vehicle only of the same volume. After 12 hours, a second dose of CL,316-243 (0.1 mg/kg) or FT895 (10 mg/kg in 5% DMA/1% Tween 80/94% sterile water) or vehicle (5% DMA/1% Tween 80/94% sterile water) was administered by i.p. injection, and after 1 subsequent hour, animals were euthanized and tissues harvested. Throughout the experiment, mice were given ad libitum access to chow and water.

All animal studies were conducted using a protocol approved by the Institutional Animal Care and Use Committee of the University of Colorado Anschutz Medical Campus, following appropriate guidelines and in accordance with the United States Public Health Service Policy on Humane Care and Use of Laboratory Animals.

#### Statistical analysis

All data are presented as mean +SEM. Statistical significance (*P* <0.05) was determined using unpaired t-test (two groups) or one-way ANOVA with correction for multiple comparisons via a Tukey post-hoc test (GraphPad Prism 9) unless otherwise specified.

## Supporting information

Expanded Materials and Methods

Full gel images

## Author contributions

E.L. Robinson, R.A. Bagchi, J. L. Major and T.A. McKinsey conceived and designed the study. E.L. Robinson, R.A. Bagchi and J.L. Major acquired data. E.L. Robinson, R.A. Bagchi, J.L. Major, J.L. Matsuda and T.A. McKinsey analyzed and interpreted data. E.L. Robinson, R.A. Bagchi, J.L. Major, B.C. Bergman, J.L. Matsuda and T.A. McKinsey wrote and reviewed the manuscript. B.C. Bergman and J.L. Matsuda provided administrative, technical, or material support. T.A. McKinsey supervised the study. The order of the first or last authors reflects the leadership exerted in the study.

## Acknowledgments

The authors wish to acknowledge the Regional Mouse Genetics Core Facility, which is supported by both National Jewish Health and the University of Colorado Anschutz Medical Campus. Metabolic phenotyping studies were performed in the Colorado Nutrition Obesity Research Center (NORC) Animal Satellite Facility (NIDDK DK048520). We thank K.M. Gavin for human SC adipocyte isolation, E. Seto for the HDAC11 antibody, J. Miano for the rat gravin-α cDNA construct, and T. Hu for cloning. E.L. Robinson was funded by the American Heart Association (Grant 829504), and R.A. Bagchi and J.L. Major received support from the Canadian Institutes of Health Research (Grants FRN-216927 and FRN-395620, respectively). T.A. McKinsey. received funding from National Institute of Health by grants HL116848, HL147558, DK119594, HL127240, HL150225, and a grant from the American Heart Association (16SFRN31400013). Contents are the authors’ sole responsibility and do not necessarily represent official NIH views.

## Supplementary Materials

- Expanded Materials & Methods
- Supplemental Figure 1. Validation of FT895 selectivity for HDAC11.
- Supplemental Figure 2. Validation of UCP1 antibody specificity.
- Supplemental Table 1. Primer sequences for genotyping, qRT-PCR and short hairpin RNA.
- Supplemental Table 2. Chemicals and reagents.
- Supplemental Table 3. Patient information for human VAT samples.
- Supplemental Table 4. Patient information for human SC adipocyte isolation.
- Supplemental Table 5. Antibodies.
- Supplemental References.

