## Supplemental Figures for "HDAC11 inhibition triggers bimodal thermogenic pathways to circumvent adipocyte catecholamine resistance"

### Supplemental Figure 1

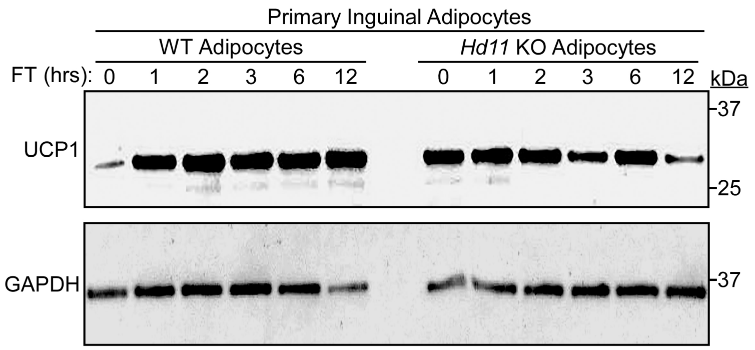


**Supplemental Figure 1. Validation of FT895 selectivity for HDAC11.** Primary adipocytes were isolated from inguinal white adipose tissue from WT and *Hdac11* global knockout (*Hd11* KO) mice and were exposed, in culture, to FT895 for the indicated times prior to homogenization and immunoblotting to assess UCP1 expression; GAPDH served as a loading control.

### Supplemental Figure 2

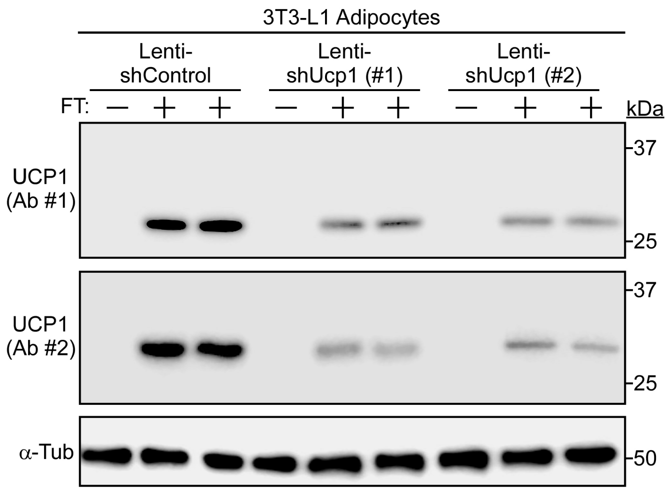


**Supplemental Figure 2. Validation of UCP1 antibody specificity.** 3T3-L1 adipocytes were infected with lentivirus encoding a control short-hairpin RNA (Lenti-shControl) or viruses encoding two independent shRNAs to target *Ucp1* mRNA (shUcp1). Cells were homogenized 24-hours after infection and immunoblot analysis was performed to assess UCP1 protein expression; α-tubulin (α-Tub) served as a loading control.
