## Supplemental Tables for "HDAC11 inhibition triggers bimodal thermogenic pathways to circumvent adipocyte catecholamine resistance"

**Supplemental Table 1. Primer sequences for genotyping and qRT-PCR**

| Primer Name | Application | Species | Primer Sequence |
|---|---|---|---|
| Ucp1 F | RT-qPCR | Human | 5'-AGTTCCTCACCGCAGGGAAAGA-3' |
| Ucp1 R | RT-qPCR | Human | 5'-GTAGCGAGGTTTGATTCCGTGG-3' |
| Ucp1 F | RT-qPCR | Mouse | 5'-CCGAAACTGTACAGCGGTCT-3' |
| Ucp1 R | RT-qPCR | Mouse | 5'-CCGAGAGAGGCAGGTGTTTC-3' |
| 18S F | RT-qPCR | Human, Mouse | 5'-GCCGCTAGAGGTGAAATTCTTA-3' |
| 18S R | RT-qPCR | Human, Mouse | 5'-CTTTCGCTCTGGTCCGTCTT-3' |
| Hdac11 flox F | DNA PCR | Mouse | 5'-GTGCAGGCCTTGGGCCTTGGCA-3' |
| Hdac11 flox R | DNA PCR | Mouse | 5'-CTGAGGAGGTAGTATGGATAG-3' |

**Supplemental Table 2. Chemicals and reagents**

| Item | Vendor | Catalog # |
|---|---|---|
| Dulbecco's Modification of Eagle's Medium (DMEM) | Corning | 10-013-CV |
| (Minimum Essential Medium α (MEMα) | Corning | 10-022-CV |
| Fetal Bovine Serum, HyClone Characterized FBS, US Origin | GE Healthcare | SH30071.02 |
| Newborn Calf Serum, heat inactivated, New Zealand origin | Thermo Scientific | 26010074 |
| Penicillin-Streptomycin-L-Glutamine, 100X | Corning | 30-009-CI |
| Insulin-Transferrin-Selenium, 100X | Thermo Gibco | 41400045 |
| Dexamethasone | Sigma Aldrich | D4902 |
| 3-Isobutyl-1-methylxanthine, IBMX | Sigma Aldrich | I5879 |
| Rosiglitazone | Cayman Chemicals | 71740 |
| Collagenase II | Worthington | LS004176 |
| Cell-permeable Cre recombinase, TAT-Cre (Tat-NLS-Cre, HTNC, HTNCre) | Excellgen | EG-1001 |
| Halt™ Protease and Phosphatase Inhibitor Cocktail | Thermo Scientific | 78440 |
| Pierce™ BCA Protein Assay Kit | Thermo Scientific | PI23227 |
| 4-15% Criterion TGX Precast Midi Protein Gel, 26 well, | BIO-RAD | 5671085 |
| 4-15% Criterion TGX Precast Midi Protein Gel, 18 well | BIO-RAD | 5671084 |
| Nitrocellulose membrane 0.45 µm | BIO-RAD | 1620115 |
| Precision Plus Protein Dual Color Standards | BIO-RAD | 1610394 |
| Bovine Serum Albumin Fraction V for immunoblotting | Akron Biotech | AAJ64655 |
| Bovine Serum Albumin, fatty acid free for immunofluorescence | Thermo Scientific | A8806 |
| Stainless steel beads 1.6 mm | Next Advance | #SSB16 |
| Alkynyl Myristic Acid (Alk-12) | Click Chemistry Tools | 1164 |

| | | |
|---|---|---|
| Click-iT® Protein Reaction Buffer Kit | Thermo Scientific | C10276 |
| Biotin Azide (PEG4 carboxamide-6-Azidohexanyl Biotin) | Thermo Scientific | B10184 |
| Pierce™ Streptavidin Magnetic Beads | Thermo Scientific | 88816 |
| FT895; FT | MedChemExpress | 2225728-57-2 |
| CL-316,243 | Sigma Aldrich | C5976 |
| Isoproterenol; ISO | Calbiochem | 420355 |
| Forskolin; FSK | Sigma Aldrich | F6886 |
| H89 | Tocris Bioscience | 2910 |
| DMSO | Sigma Aldrich | D8418 |
| N,N-Dimethylacetamide (DMA) | Sigma Aldrich | D137510 |
| Tween-80 | Sigma Aldrich | P1754 |
| 4,4-Difluoro-1,3,5,7,8-Pentamethyl-4-Bora-3a,4a-Diaza-s-Indacene; BODIPY | Thermo Scientific | D3922 |
| 4',6-diamidino-2-phenylindole; DAPI | Thermo Scientific | D1306 |
| D-Biotin | Sigma Aldrich | B4639 |
| D-Pantothenic acid hemicalcium salt | Sigma Aldrich | P5155 |
| 3,3′,5-Triiodo-L-thyronine sodium salt; T3 | Sigma Aldrich | T6397 |
| Transferrin, human | Sigma Aldrich | T8158 |
| Insulin solution, human | Sigma Aldrich | I9278 |
| Tween-20 | Sigma Aldrich | P1754 |
| Glass coverslips, 25 mm | Chemglass | CLS-1763-025 |
| Microscope slides, Diamond White Glass, 25 x 75mm, Charged, 90° Ground Edges, White Frosted | Global Scientific | 1358W |
| Nunc™ Lab-Tek™ II Chamber Slide™ | Thermo Scientific | 12-565-7 |
| Formaldehyde,16 % methanol-free Ultra Pure | Thermo Scientific | 18814-10 |
| VECTASHIELD® HardSet™ Antifade Mounting Medium for immunofluorescence | Vector Labs | H-1400-10 |
| Antigen Unmasking Solution, Citrate-Based | Vector Labs | H-3300-250 |
| BLOXALL® Endogenous Blocking Solution, Peroxidase and Alkaline Phosphatase | Vector Labs | SP-6000-100 |
| ECTASTAIN® Elite® ABC-HRP Kit, Peroxidase (Standard) | Vector Labs | PK-6100 |
| Vector® NovaRED™ Substrate Kit, Peroxidase (HRP) | Vector Labs | SK-4800 |
| Fisher Chemical™ Permount™ Mounting Medium for IHC | Thermo Scientific | SP15-100 |
| Agarose | Life Science Products | A-1705 |
| 1kb DNA ladder | Promega | PR-G5711 |
| Protein ladder, Precision Plus Protein™ Dual Color Standards | BIO-RAD | 1610394 |
| Non-fat milk powder, Kroger® Instant Non-Fat Dry Milk | Kroger® | 0001111083297 |
| PowerUp SYBR Green Master Mix | Thermo Scientific | 4368577 |

| | | |
|---|---|---|
| TRIzol™ Reagent | Thermo Scientific, Ambion | 15596026 |
| Verso cDNA Synthesis Kit | Thermo Scientific | AB1453B |
| Polybrene | Sigma Aldrich | TR-1003 |
| Polyethylenimine, Linear, PEI | Polysciences Inc. | 23966-1 |

**Supplemental Table 3. Patient information for human VAT samples**

| Age (yrs) | Sex | Ethnicity | BMI Consent | BMI Surgery |
|---|---|---|---|---|
| 30 | F | Hispanic | 42.0 | 42.0085992 |
| 39 | M | Non-Hispanic, Indian | 36.9839009 | 36.9839009 |
| 22 | F | White | 42.0 | 41.8102305 |

**Supplemental Table 4. Patient information for human SC adipocyte isolation**

| Age (yrs) | Sex | BMI |
|---|---|---|
| 29 | M | 21.1 |

**Supplemental Table 5. Antibodies**

| Antigen | Vendor | Host | Catalog # |
|---|---|---|---|
| UCP1 #1 | Abcam | Rabbit | ab209483 |
| UCP1 #2 | Cell Signaling Technology | Rabbit | 14670S |
| GAPDH | Thermo Scientific | Mouse | AM4300 |
| Phospho-PKA substrates | Cell Signaling Technology | Rabbit | 9624 |
| $\alpha$-Tubulin-HRP | Santa Cruz Biotechnology | Mouse | sc-23948 |
| Gravin-α | Proteintech | Rabbit | 25199-1-AP |
| FLAG-HRP | Sigma Aldrich | Mouse | F1804 |
| $\beta_3$-AR | Abcam | Rabbit | ab94506 |
| $\beta_2$-AR | Proteintech | Rabbit | 13096-1-AP |
| HDAC11 | From Dr. Edward Seto, George Washington University | Rabbit | |
| anti-Rabbit Alexa Fluor 594 for Immunofluorescence | Thermo Scientific | Goat | A-11012 |
| Anti-Rabbit IgG-HRP | Southern Biotech | Goat | OB4050-05 |
| Anti-Mouse IgG-HRP | Southern Biotech | Goat | OB1031-05 |
| Anti-Rabbit IgG Antibody (H+L), Biotinylated for IHC | Vector Labs | Goat | BA-1000-1.5 |
