## Supplementary material for "HDAC11 inhibition triggers bimodal thermogenic pathways to circumvent adipocyte catecholamine resistance": Expanded Materials and Methods

### Supplemental Material

### Expanded Materials and Methods

*Human visceral adipose tissue (VAT) explant acquisition and treatment.* Human VAT samples were acquired from bariatric surgery patients under protocols approved by Colorado Multiple Institutional Review Board. All VAT samples were deidentified prior to use in experiments. Patient characteristics are described in Supplemental Table 3. VAT explants were washed, trimmed and rinsed in pre-warmed DMEM and cut with fine scissors into 1 mm x 1 mm pieces for culture in ThinCert™ Tissue Culture Inserts (Greiner Bio-One) submerged in 1 mL DMEM in 6-well tissue culture plates. Human VAT explants were treated with FT895 (100 μM), CL-316,243 (10 μM), isoproterenol (100 μM) or forskolin (100 μM) or vehicle control (DMSO; 0.1% final concentration). VAT explants were cultured and maintained in a humidified cell culture incubator maintained at 37°C with 5% $CO_2$.

*Human subcutaneous (SC) preadipocyte acquisition, culture and differentiation*. Human SC adipose tissue samples were acquired from laparoscopic surgeries from volunteers, as described previously, described under protocols approved by Colorado Multiple Institutional Review Board (NCT02654925) (1). Subject information can be found in Supplemental Table 4. Stromal vascular fraction (SVF)-derived preadipocytes were isolated according to published procedures and plated onto 6-well tissue culture dishes in growth media (MEMα, 10% FBS, 0.1X Penicillin-Streptomycin-Glutamine) (2). Once

SVF-derived preadipocytes reached 90% confluency, medium was changed to complete differentiation induction media for up to seven days, with medium refreshed every 48 hours (DMEM, 1X PSG, 33 μM d-Biotin, 17 μM pantothenate, 100 nM dexamethasone, 1μM rosiglitazone, 0.5 mM IBMX, 2nM T3, 10 μg/mL transferrin, 100 nM insulin). Following seven days, complete differentiation medium was switched to maintenance media (DMEM, 1X PSG, 33 μM d-Biotin, 17 μM pantothenate, 10 nM dexamethasone, 100 nM insulin). Differentiation was assessed by lipid droplet coverage by brightfield microscopy (EVOS FL, Life Technologies), with 80-90% lipid droplet coverage considered fully differentiated.

*Adipocyte culture.* The 3T3L1 CL-173™ preadipocyte cell line was purchased from ATCC. Cells were maintained and differentiated into mature adipocytes *in vitro* as previously described (3). Undifferentiated cells were seeded on culture dishes and grown to 80-90 % confluency in DMEM supplemented with 10% newborn calf serum and 1X Penicillin-Streptomycin-L-Glutamine. Differentiation of 3T3-L1 cells was induced using DMEM supplemented with 10% fetal bovine serum, 1 μM dexamethasone, 0.5 mM 3-isobutyl-1-methylxanthine, 1 μg/mL insulin for 72 hr. Following this incubation, media was changed to adipocyte maintenance media (DMEM with 1μg/mL insulin, 10% FBS) and replenished every 48h for 8 days or until lipid droplet coverage is 80 – 90 %, as assessed by brightfield microscopy. At this time, 3T3-L1 adipocytes were treated as per the experiments described.

HIB1B brown-like preadipocytes were cultured in DMEM supplemented with 10% FBS and 1X Penicillin-Streptomycin-L-Glutamine. HIB1B adipocytes were differentiated into mature adipocytes at 70% confluency using rosiglitazone (1 μM) for four days. Differentiation was monitored by assessing lipid droplet coverage by brightfield microscopy, as well as high basal UCP1 expression.

Mouse primary preadipocytes were isolated from SVF of murine inguinal AT of 10 – 12-week-old male WT, *Hdac11*$^{fl/fl}$ or *Hdac11*$^{KO}$ C57BL/6J mice, as previously described (3). ingWAT was dissected and washed in pre-warmed DMEM before being transferred to a petri dish containing 10 mL of 0.2% (w/v) collagenase II. The samples were subsequently cut into ~1 mm pieces and further mincing was performed to allow opening up of the tissue using spring scissors (Fine Science Tools). The minced tissue in digestion buffer was then transferred to a 15 mL Falcon tube with the lid loosely fastened and incubated in a shaker at 37 ºC at 400 rpm for 20 minutes, with vortexing every 5 minutes. At the end of the digestion, the tissue slurry was passed through a 70 μm cell strainer into a 50 mL Falcon tube, followed by 30 mL preadipocyte media (5% NBCS, 5% FBS,1% PSG) to stop enzymatic activity. After a brief (2-3 minute) centrifugation at 200g at room temperature, the upper floating mature adipocyte-containing layer was discarded. The cell suspension was then centrifuged at 2000 x g for 5 minutes at room temperature to pellet the SVF-derived preadipocytes. The supernatant was removed and the remaining cell pellet resuspended in pre-warmed preadipocyte media. Cells were plated directly onto 6- well cell culture plates (VWR). Once 90 - 100 % preadipocyte confluency was reached, differentiation was induced by treatment with differentiation induction medium (DMEM, 10% FBS, 0.5% PSG, 1 μg/mL insulin, 0.5 μM dexamethasone, 0.25 mM 3-isobutyl-1- IBMX and 1 um rosiglitazone) for 72 hours. On day 3, induction medium was replaced with maintenance media (DMEM, 10 % FBS, 0.5% PSG, 1 ug/mL insulin).

Maintenance media was replenished every 48 hours and maturation followed by observing lipid droplet coverage by brightfield microscopy, with 80 -90 % lipid droplet coverage considered fully differentiated.

*Plasmids and lentivirus generation.* p3XFLAG-CMV-14 expression vector encoding full length rat gravin-α was a gift from Dr. Joseph Miano, Augusta University, U.S.A (4). This base vector was used to generate the K1502/K1505 mutant construct of gravin-α, as previously described (3). These constructs were sub-cloned into pLenti CMV Hygro DEST (Addgene #17454). To create short hairpin RNA encoding lentivirus, pLKO.1 plasmids (sequences obtained from the Sigma MISSION™ shRNA library) encoding shRNA for mouse *gravin-α* (TRCN0000088805), *Ucp1* (TRCN0000114241 and TRCN0000114245) and a scrambled shRNA control (SHC002) were obtained through the Functional Genomics Facility at the University of Colorado Cancer Center. Lentiviruses were generated by co-transfection of shRNA in pLKO.1 or pLenti overexpression plasmids, psPAX2 and pMD2.G into 293T cells (ATCC #CRL-3216™). The viral supernatant was collected 60 hours after transfection, filtered through a 0.45 μm syringe filter. Viral supernatant was supplemented with polybrene at 1 μl/mL (stock concentration 10mg/mL) and used to directly infect adipocytes. After 24 or 48 hours of infection, cells were used for experiments as indicated. psPAX2 and pMD2.G were gifts from Didier Trono (Addgene plasmids # 12260, #12259).

*Assessment of gravin-α myristoylation*. Detection of lysine myristoyation on gravin-α was performed using a Click chemistry approach as previously described (3). Cells were supplemented with 50 μM alkynyl myristic acid for 4 hours prior to harvest, then lysed in 1% SDS lysis buffer (50 mM Tris-HCl pH 8.0, 1% (w/v) SDS) with protease and phosphatase inhibitors. Cell lysates were prepared as standard and protein concentration was determined using a BCA Protein Assay Kit. Click chemistry reactions with biotin-azide (50 μM final concentration) were performed at room temperature for 30 minutes with end-over-end rotation with up to 200 μg of protein per sample, and the reactions were terminated by bringing the volume of the sample to 60 μl with 1% SDS lysis buffer. To precipitate and concentrate proteins, 600 μL of methanol, 150 μL of chloroform and 400 μL of 18 MΩ water were added to each sample. Samples were vortexed and centrifuged at 13,000 x g for 5 minutes at room temperature. The upper aqueous supernatant was gently removed and the interface layer containing the protein precipitate was left intact. 450 μL of methanol was added to the tubes, vortexed, and the samples spun at 13,000 x g for 5 minutes to pellet the protein. The methanol supernatant was removed, and the protein pellets were washed once again with 450 μL of methanol or until the protein pellets are white in color. The protein pellets were then air-dried for 15-30 minutes and resolubilized in 1% SDS buffer. Protein concentration was assessed by BCA assay, and 100 μg of biotin-labeled protein was then incubated with 50 μL of streptavidin magnetic beads overnight at 4°C with end-over-end rotation. Following incubation, beads were collected with a magnetic rack and rinsed three times with washing buffer (Tris-buffered saline containing 0.1% Tween™-20). After the final wash, the supernatant was discarded and 50 uL 2X sample buffer (100 mM Tris pH 6.8, 4% SDS w/v, 20% glycerol v/v, 10% β-mercaptoethanol and bromophenol blue) was added to the beads and boiled at 95 °C for 10 minutes.  The samples were further analyzed via immunoblotting using antibody specific for gravin-$\alpha$, and total gravin-$\alpha$ amount was used for reference input.

*Immunoblotting.* Cultured cells were washed with PBS and lysed with RIPA buffer (50mM Tris.HCl pH 8.0, 150mM NaCl, 1% NP-40, 0.5% sodium deoxycholate and 0.1% SDS supplemented with protease and phosphatase inhibitors) for 30 minutes on ice followed by scraping and centrifugation at 13,000 xg for 5 minutes. For AT, RIPA buffer was added to 25-50 mg of tissue in an Eppendorf tube and the tissue was physically disrupted with sterile stainless steel beads in a bullet blender (Next Advance Bullet Blender 24) for 5 minutes at setting #9 at 4 ºC. Protein concentrations were measured using a BCA assay, and 10 – 30 $\mu$g of protein was resolved through 4-15 % pre-cast polyacrylamide gels and transferred onto 0.45 $\mu$m nitrocellulose membranes. Membranes were blocked in 5% non-fat milk powder in TBS-Tween (20 mM Tris, 150 mM NaCl, 0.1% Tween 20) and proteins were analysed by immunoblotting using antibodies against specific antigens (Supplemental Table 5). Membranes were incubated with primary antibodies at a final concentration of 1:1000 in 2.5% bovine serum albumin in TBS-Tween for 2 hours at room temperature or overnight at 4 ºC on a rocker. Washes were performed thrice for 10 minutes in TBS-Tween followed by secondary antibody incubation with horse radish peroxidase (HRP)-conjugated anti-rabbit or anti-mouse secondary antibodies at a final concentration of 1:2000 in 5% non-fat milk in TBS-Tween. Final washes were performed in 1 X TBS-Tween, thrice for 10 minutes, prior to imaging. Immunoblot images were acquired on an Odyssey® XF digital imaging system (LI-COR) using the chemiluminescence program and LI-COR Acquisition software or on a ChemiDoc™

Imaging system (BIO-RAD). Protein abundance was quantified using the LI-COR Image Studio™ Lite software and normalized to respective reference proteins. Immunoblotting with homogenates of 3T3-L1 cells that were infected with two different lentiviral constructs expressing short hairpin RNA sequences against *Ucp1* confirmed the specificity of the anti-UCP1 antibodies that were employed (Supplemental Figure 2).

*Reverse transcription Quantitative PCR (RT-qPCR) gene expression analysis.* RNA was isolated from cells or tissue using TRIzol lysis reagent and 0.5 – 1.0 μg of RNA was reverse transcribed and cDNA synthesized with random hexamers for single strand synthesis. RT-qPCR was performed on a StepOnePlus Real-Time PCR System (Applied Biosystems) using SYBR Green Master Mix and gene-specific primers at a final concentration of 500 nM each for forward and reverse primers. RT-qPCR primer sequences are provided in Table 1. Relative gene expression was calculated using the $2^{-ddCt\Delta CT}$ method and normalized to 18S RNA (5).

*Immunofluorescence microscopy.* 3T3L1 cells were grown and differentiated on glass coverslips (25 mm diameter) in 6-well tissue culture plates. Cells were fixed with 4% formaldehyde in PBS for at least 20 minutes at room temperature followed by permeabilization in 0.25% Triton X-100 in PBS for 15 minutes. Non-specific antigen binding was blocked by incubating the cells with 5 % bovine serum albumin in PBS for 1 hour at room temperature. Immunostaining was performed by incubating UCP1 primary antibody (Abcam, ab209483) at a concentration of 1:250 in 2.5% BSA in PBS overnight at 4 ºC on a rocker. Cells were washed thrice with PBS and incubated with fluorescent secondary antibody (1:500) along with BODIPY (5 mM) and 4',6-diamidino-2-phenylindole (DAPI) at a final concentration of 300 nM in PBS for one hour at room temperature, rocking gently.  Cells were then washed twice in PBS then mounted onto white frosted microscope slides. Imaging was performed on a Zeiss LSM780 confocal microscope (Advanced Light Microscopy Core, University of Colorado Anschutz Medical Campus) using a 100x objective lens, and images processed through ZEN Black software (Zeiss).

For wholemount tissue imaging, AT was fixed in 4% formaldehyde in PBS overnight at 4 ºC with end-over-end rotation followed by permeabilization in 0.5% Triton X-100 in PBS for 1 hour with end-over-end rotation. Lipid droplets were stained with BODIPY (5 μM) in PBS overnight at 4 ºC with end-over-end rotation. Stained AT was rinsed twice in PBS, mounted in chamber slides and whole mount imaging was performed on a Zeiss LSM780 confocal microscope (Advanced Light Microscopy Core, University of Colorado Anschutz Medical Campus) using a 10x objective lens and processed through ZEN Black software (Zeiss).

*UCP1 immunohistochemistry.* Epididymal AT from chow fed control (*Hdac11*$^{fl/fl}$) and *Hdac11*$^{cKO}$ was formalin fixed and embedded in paraffin. Tissue sections (5 μm thick) of eWAT were deparaffinized, hydrated, and subjected to antigen retrieval using citrate buffer. Endogenous peroxidase activity was inhibited using blocking for 10 minutes, and sections were subsequently incubated in 5% BSA for 1 hour at room temperature. Tissue sections were incubated overnight with primary anti-UCP1 antibody (Abcam, ab209483) at a final concentration of 1:100, followed by incubation with biotinylated secondary antibody at a final concentration of 1:500, and immunoperoxidase detection was performed. Images were acquired using an Olympus microscope (BX51) using a 10x objective lens equipped with a multicolor camera (DP72).
