## Supplementary material for "HDAC11 inhibition triggers bimodal thermogenic pathways to circumvent adipocyte catecholamine resistance": Full gel images

### Slide 1
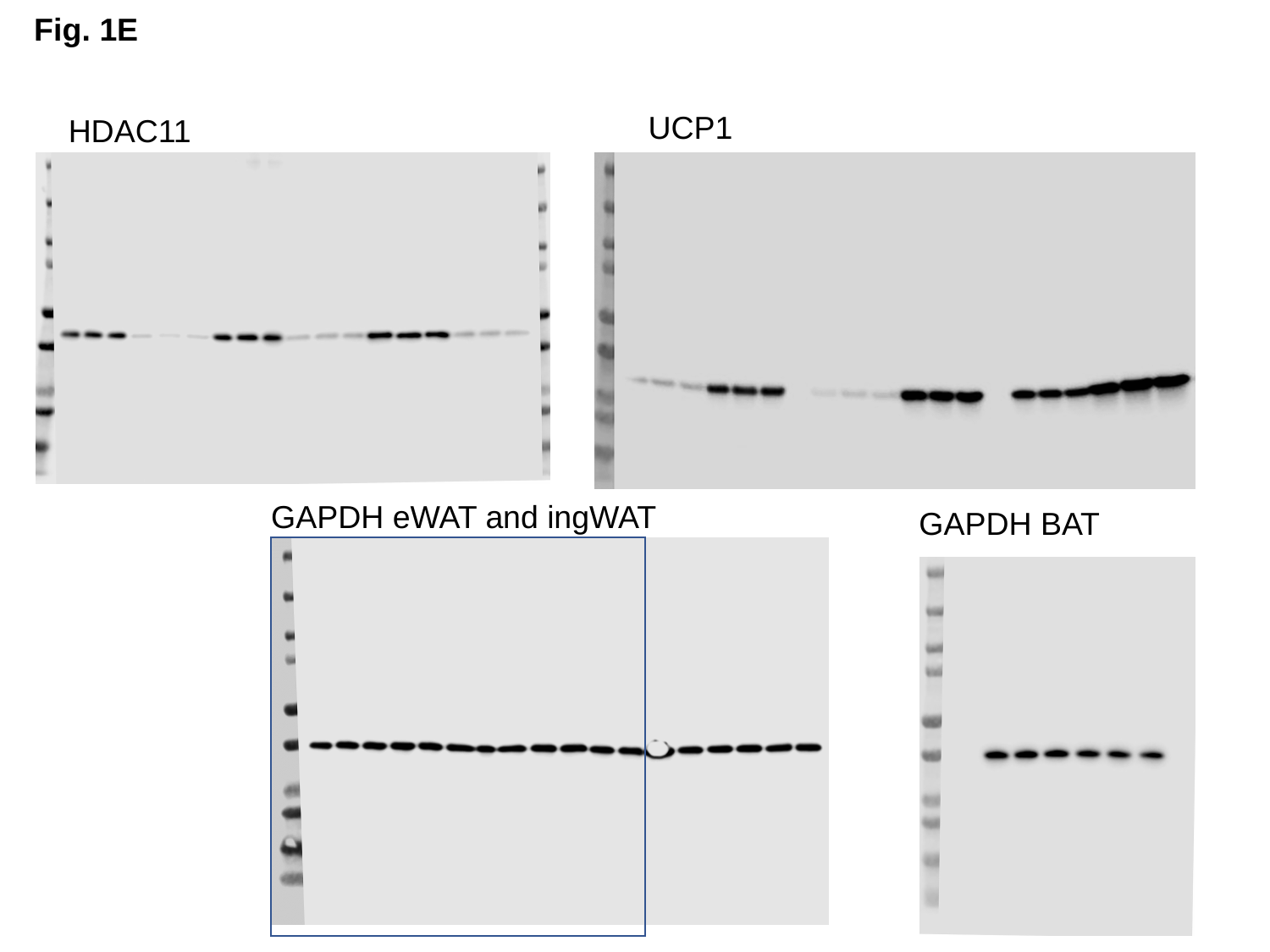

Fig. 1E
UCP1
HDAC11
GAPDH eWAT and ingWAT
GAPDH BAT

### Slide 2
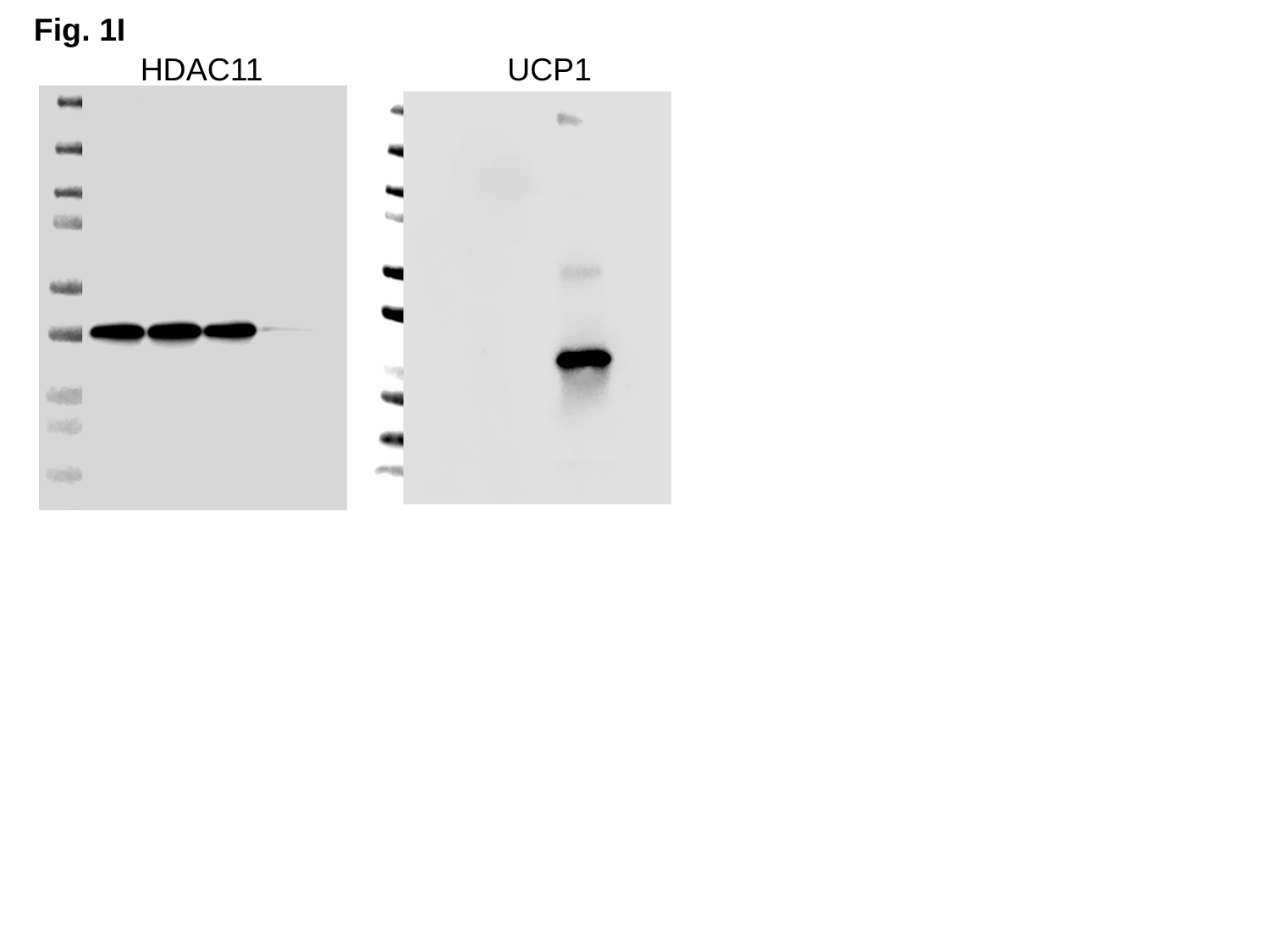

Fig. 1I
HDAC11
UCP1

### Slide 3
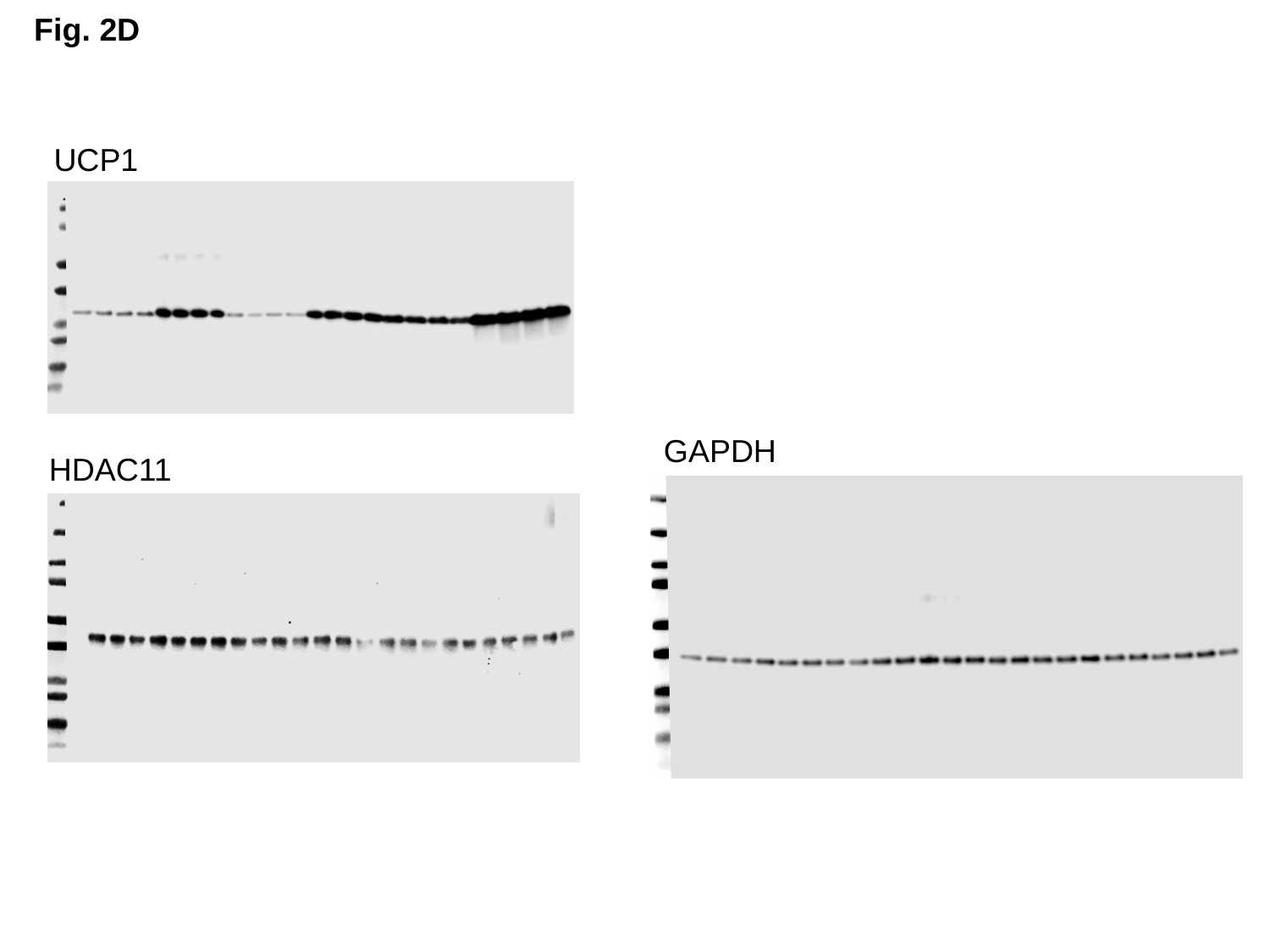

Fig. 2D
UCP1
GAPDH
HDAC11

### Slide 4
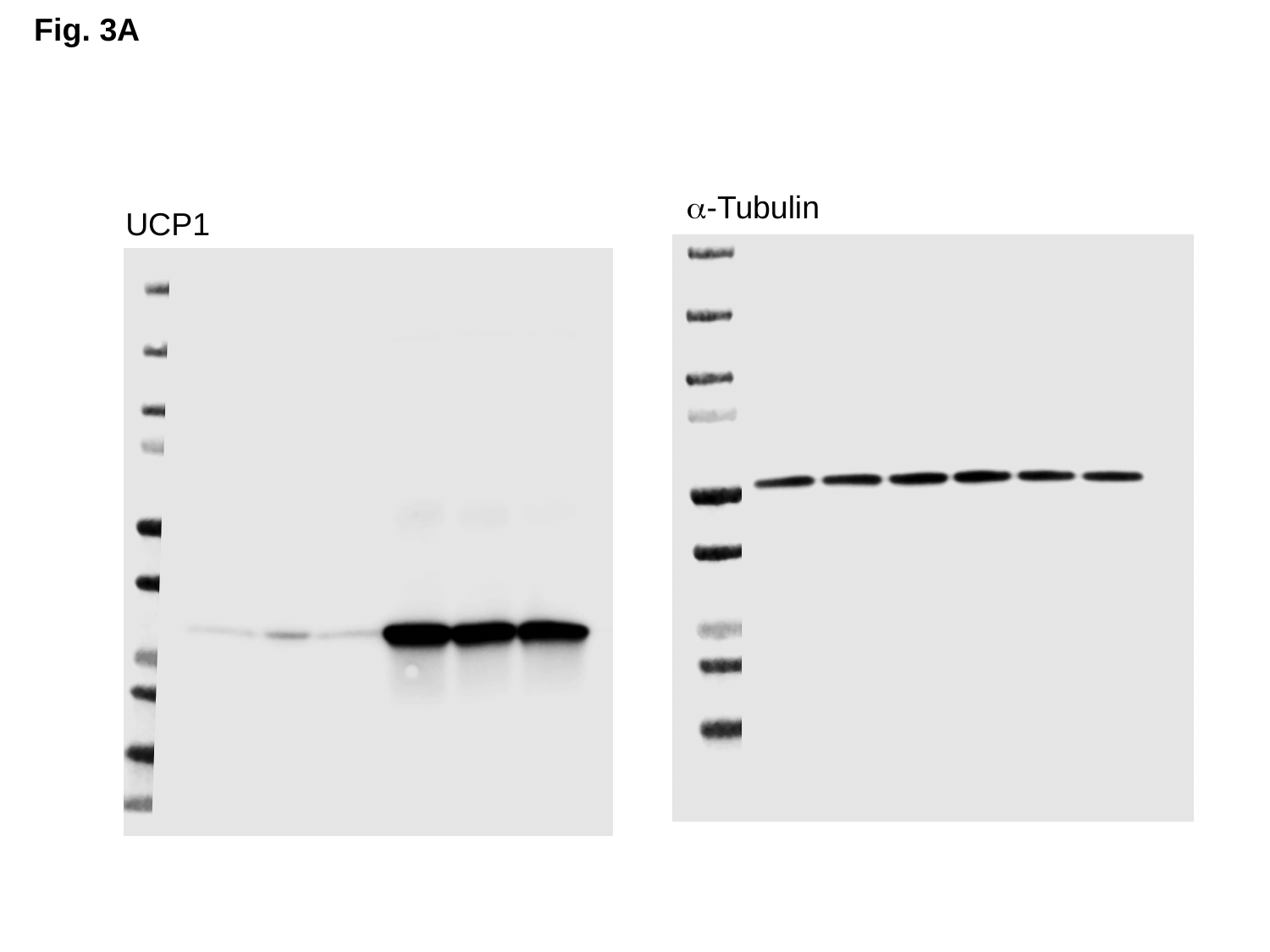

Fig. 3A
α-Tubulin
UCP1

### Slide 5
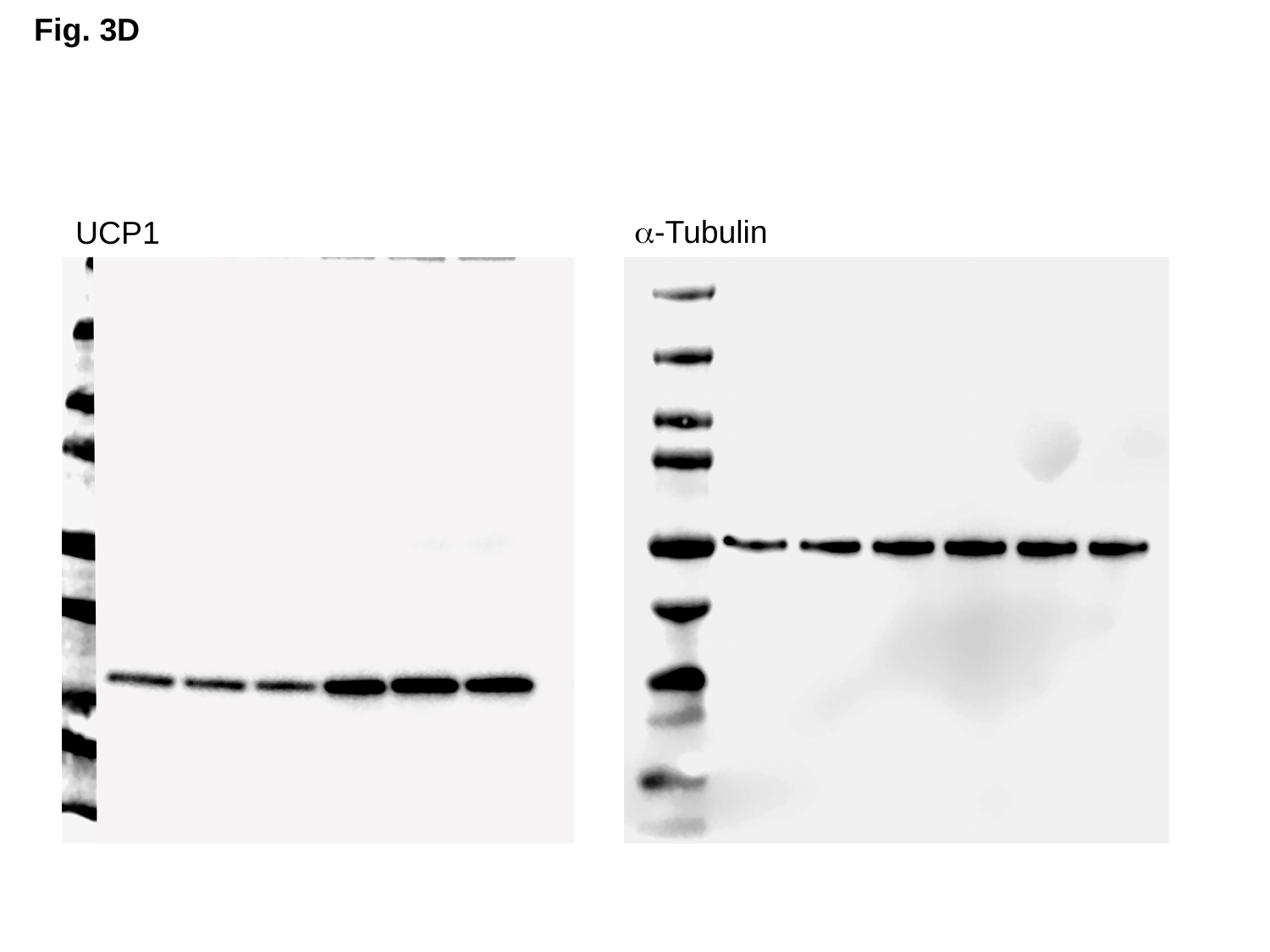

Fig. 3D
α-Tubulin
UCP1

### Slide 6
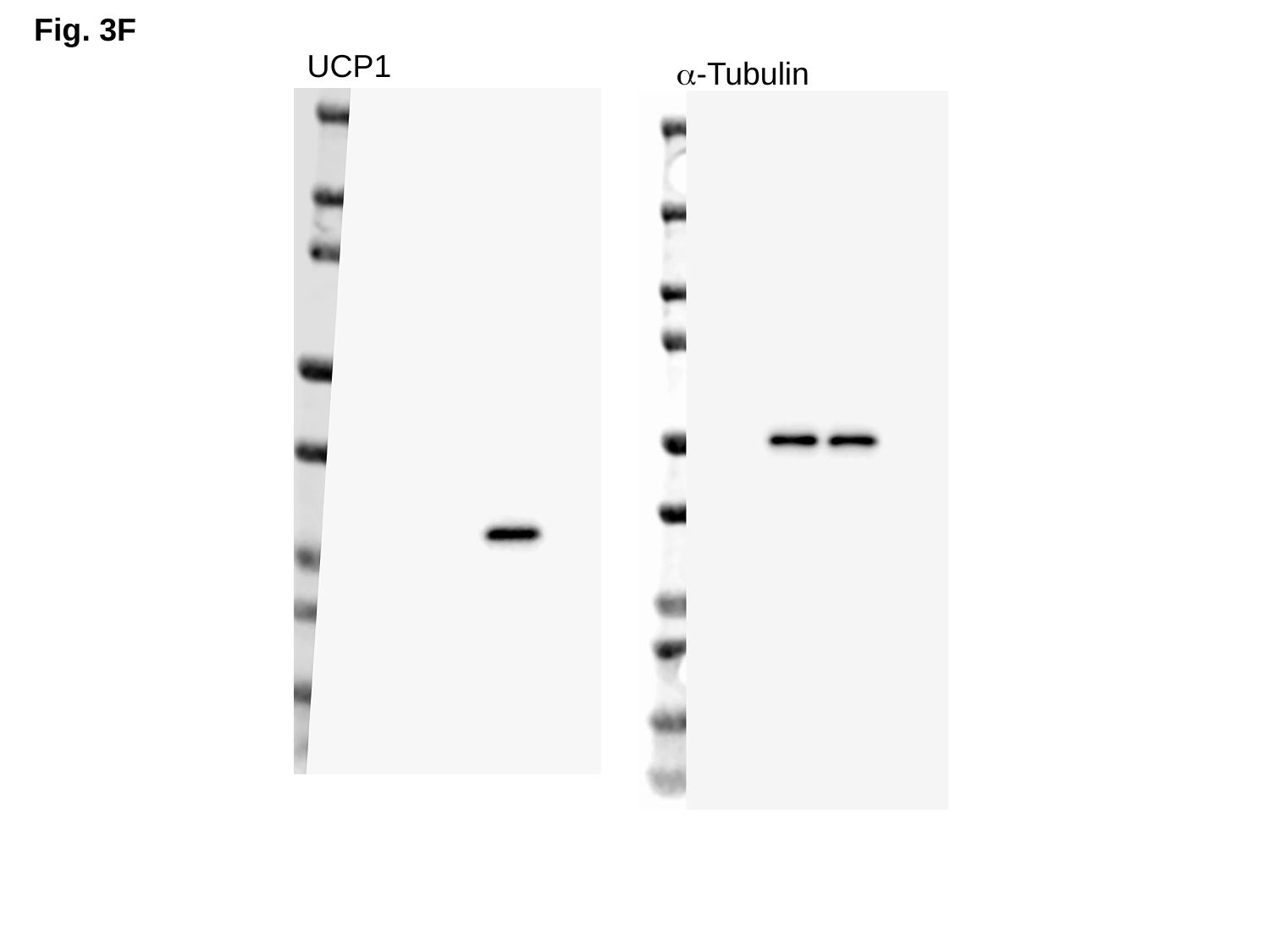

Fig. 3F
UCP1
α-Tubulin

### Slide 7
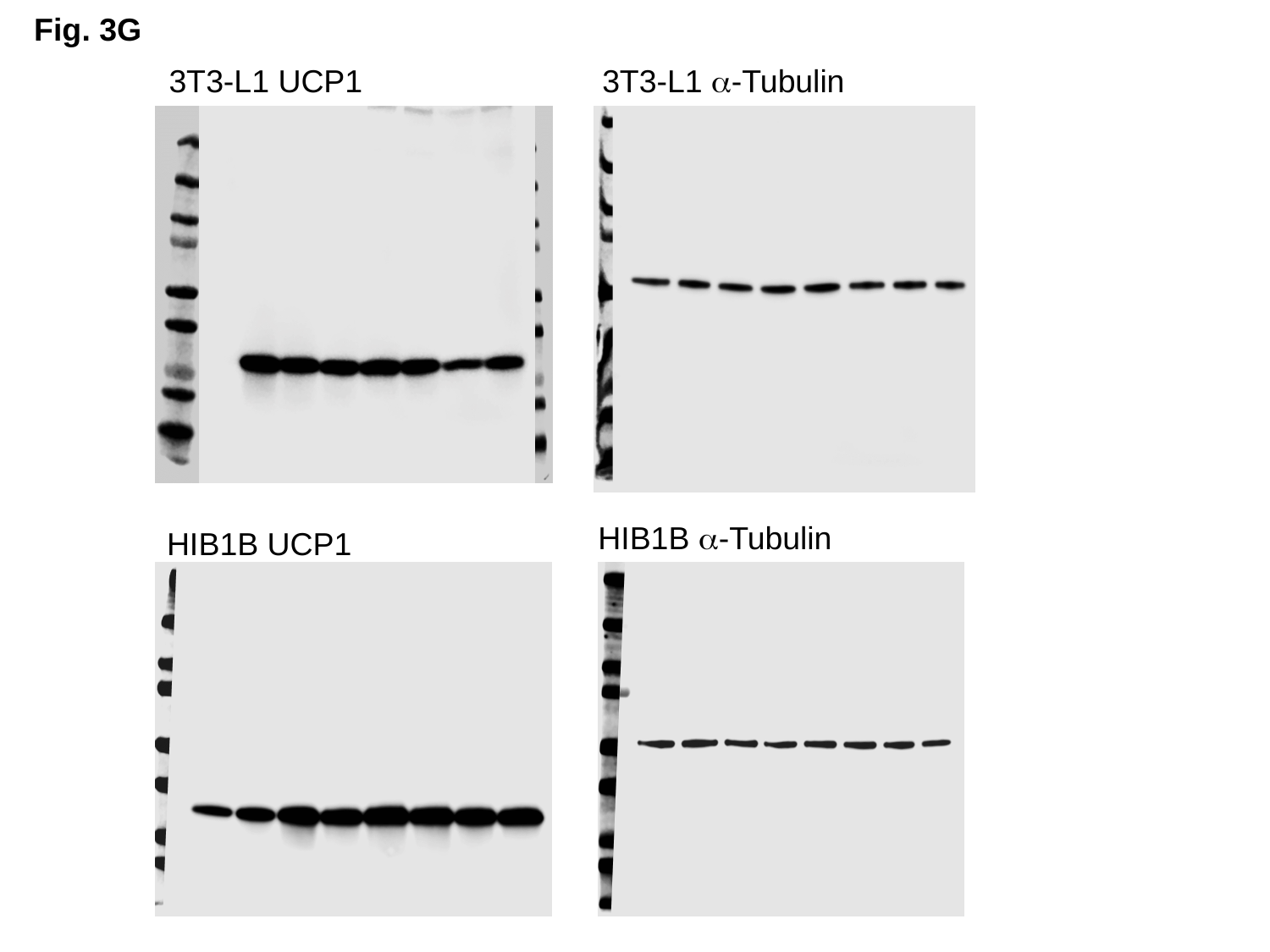

Fig. 3G
3T3-L1 UCP1
3T3-L1 α-Tubulin
HIB1B α-Tubulin
HIB1B UCP1

### Slide 8
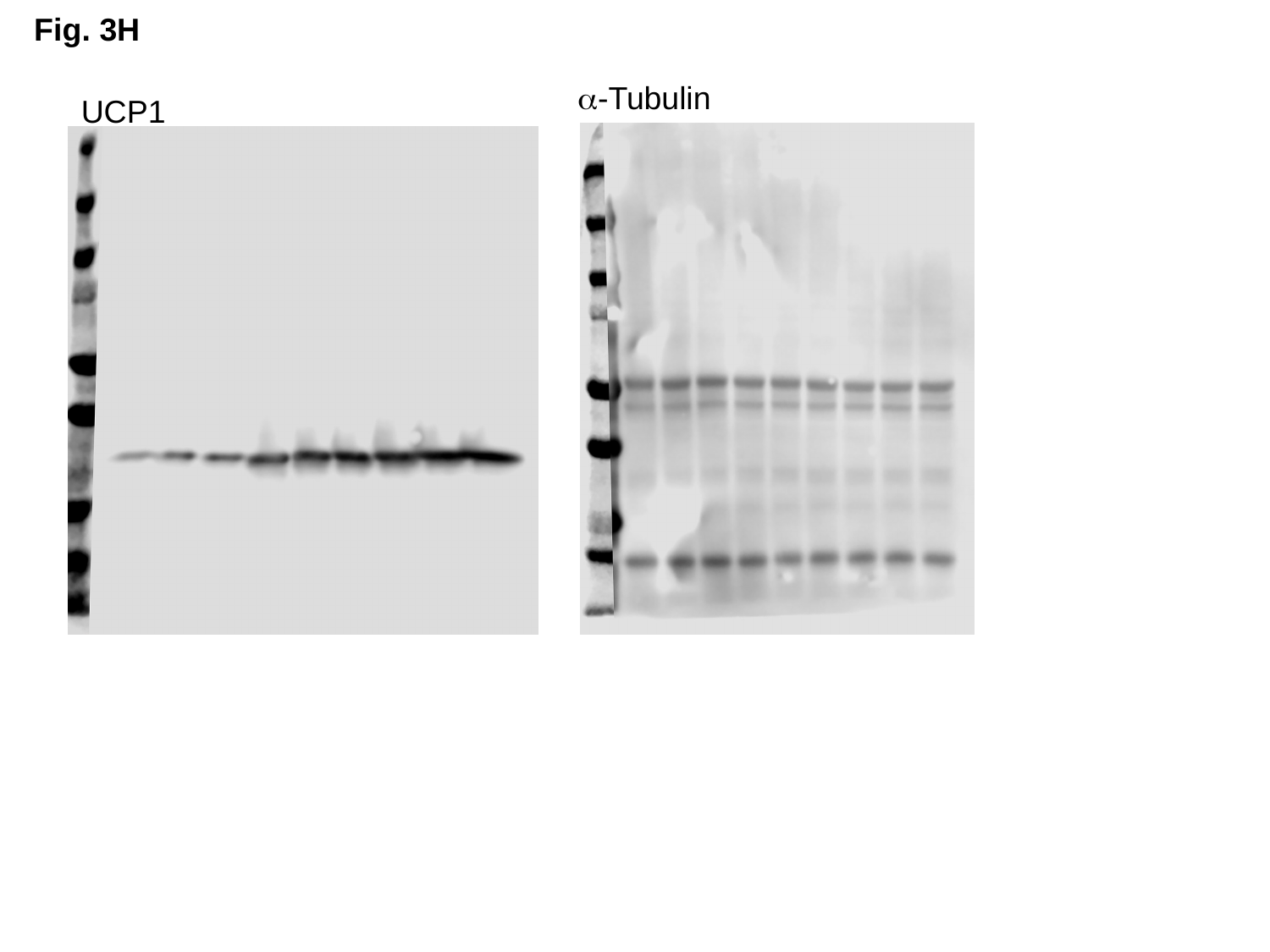

Fig. 3H
α-Tubulin
UCP1

### Slide 9
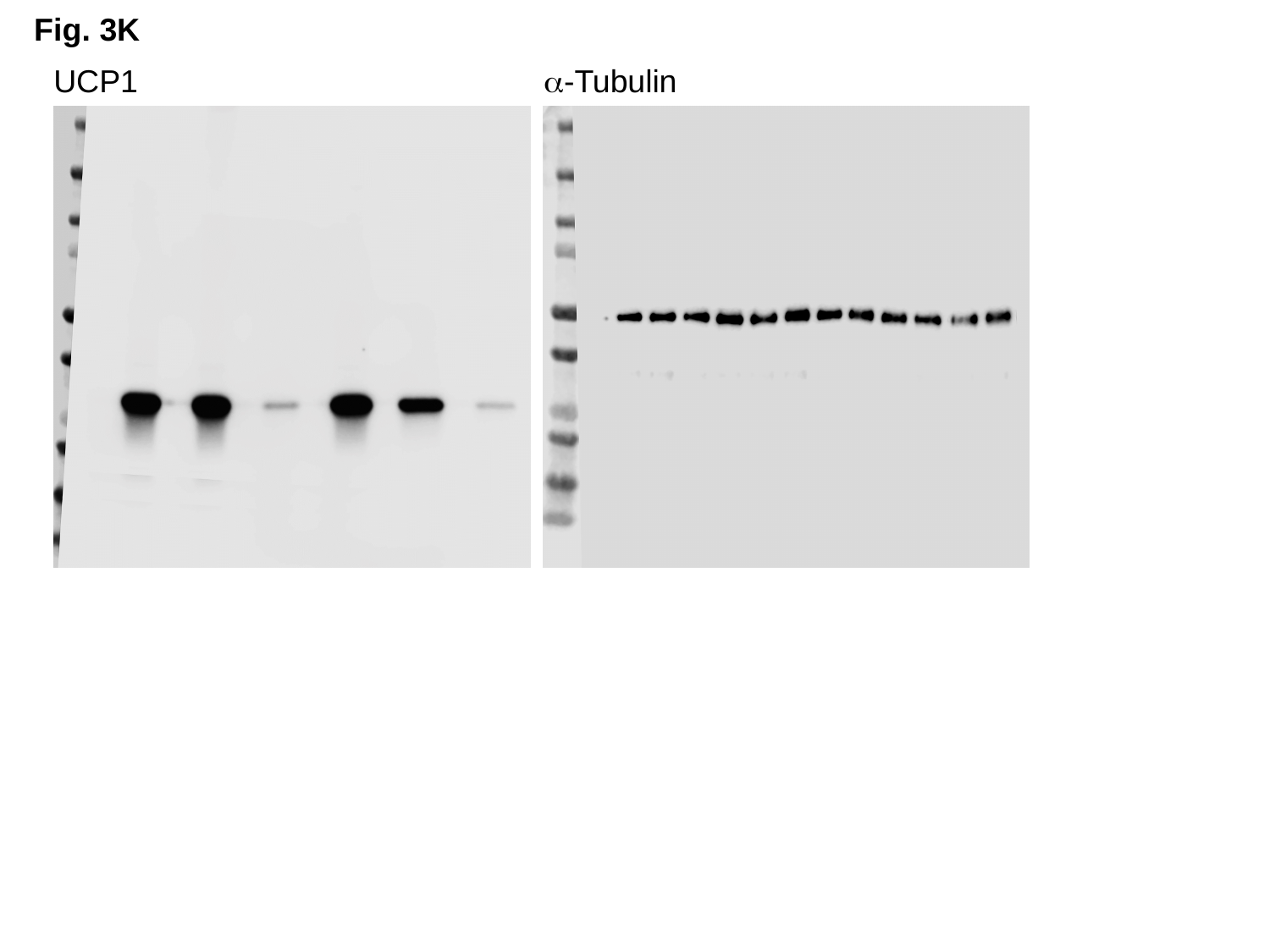

Fig. 3K
UCP1
α-Tubulin

### Slide 10
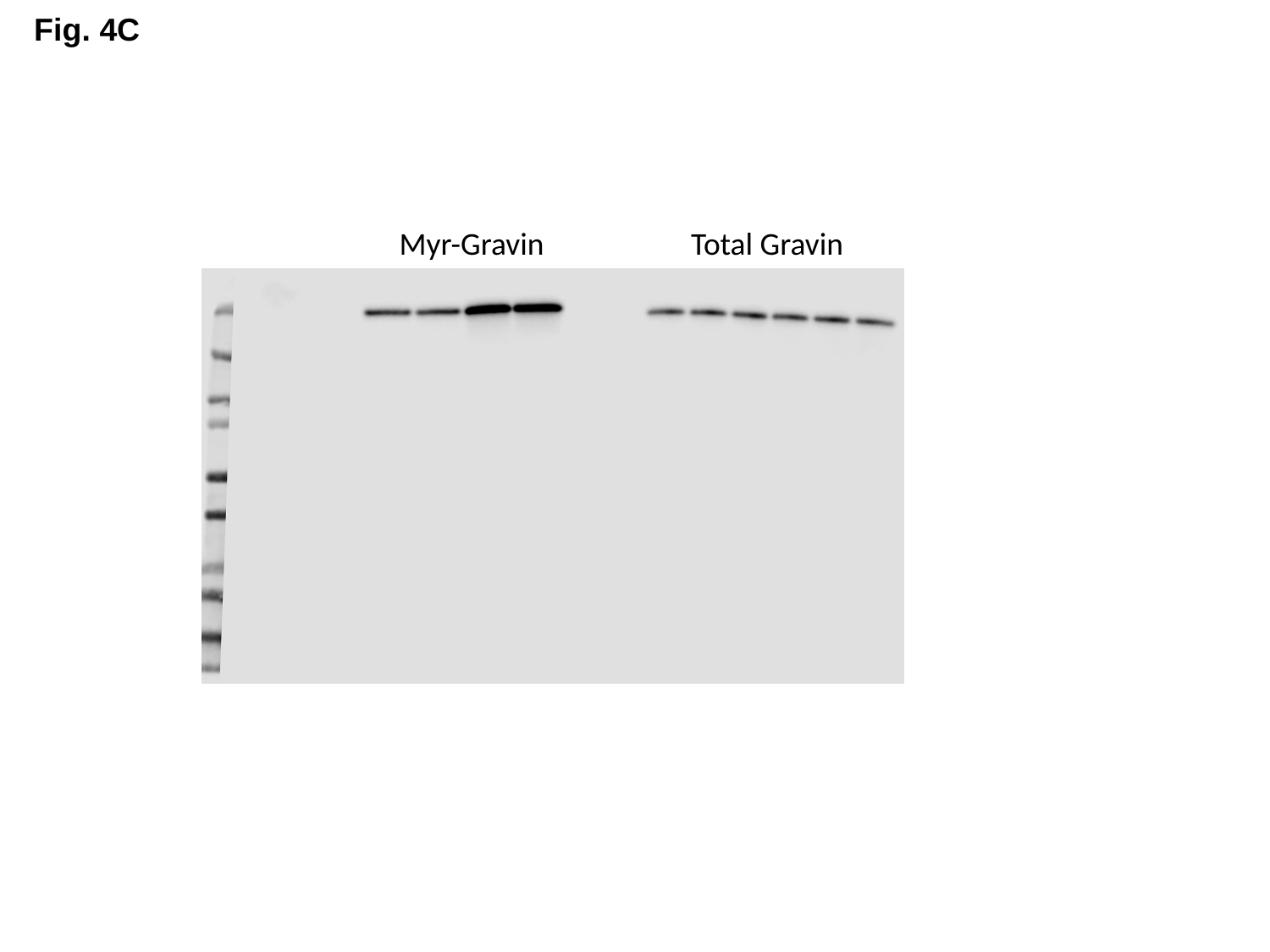

Fig. 4C
Myr-Gravin
Total Gravin

### Slide 11
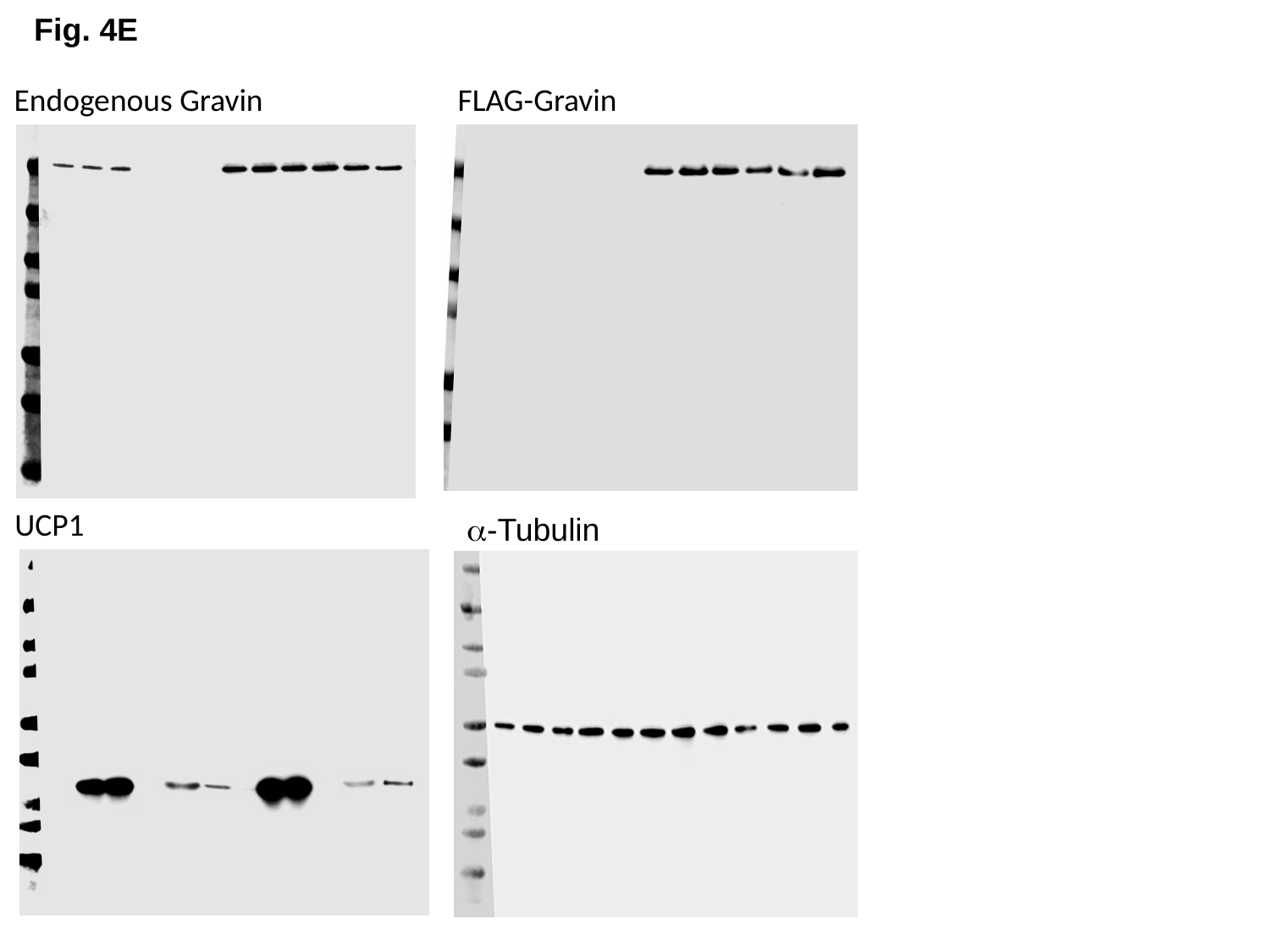

Fig. 4E
Endogenous Gravin
FLAG-Gravin
UCP1
α-Tubulin

### Slide 12
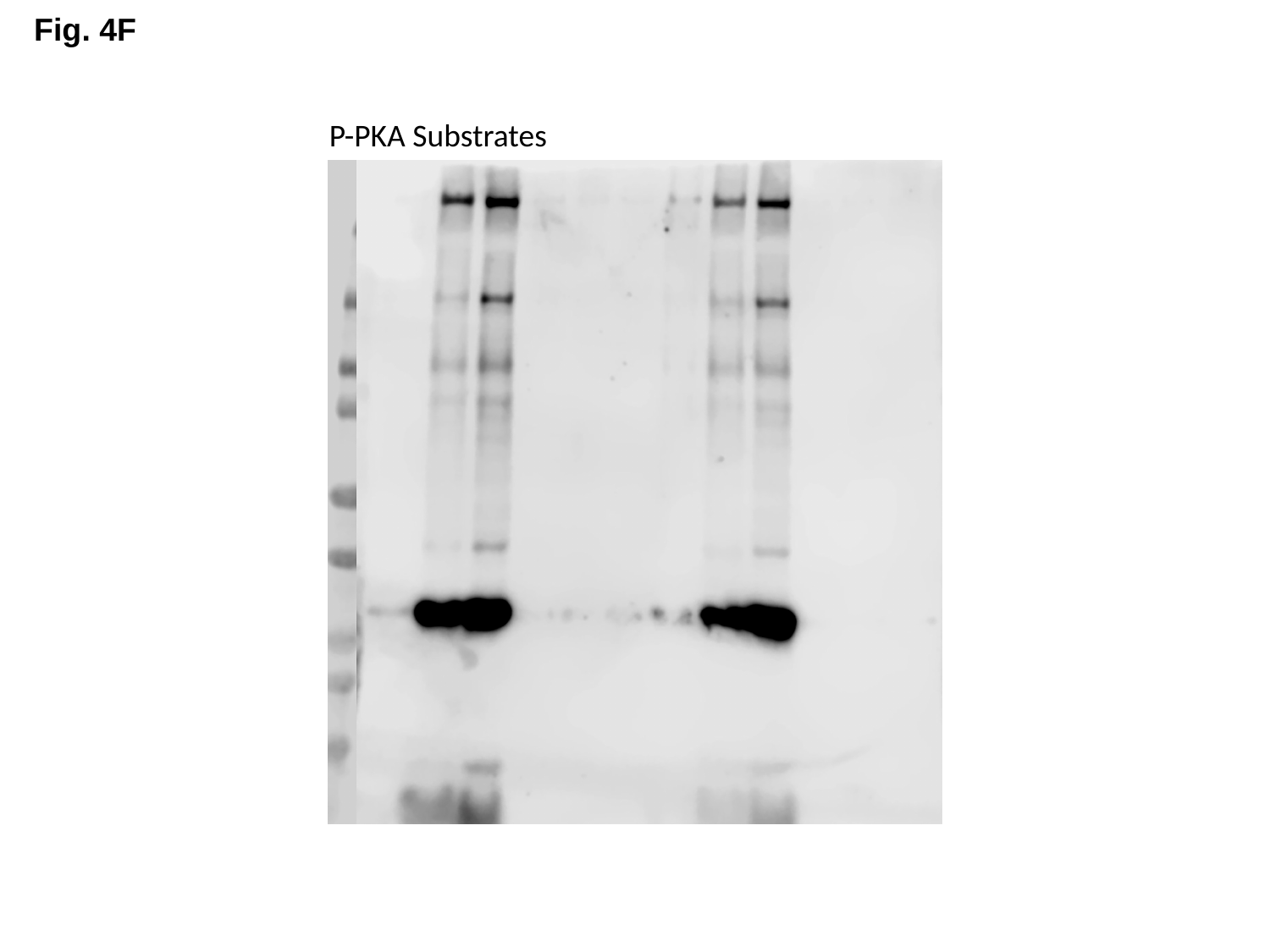

Fig. 4F
P-PKA Substrates

### Slide 13
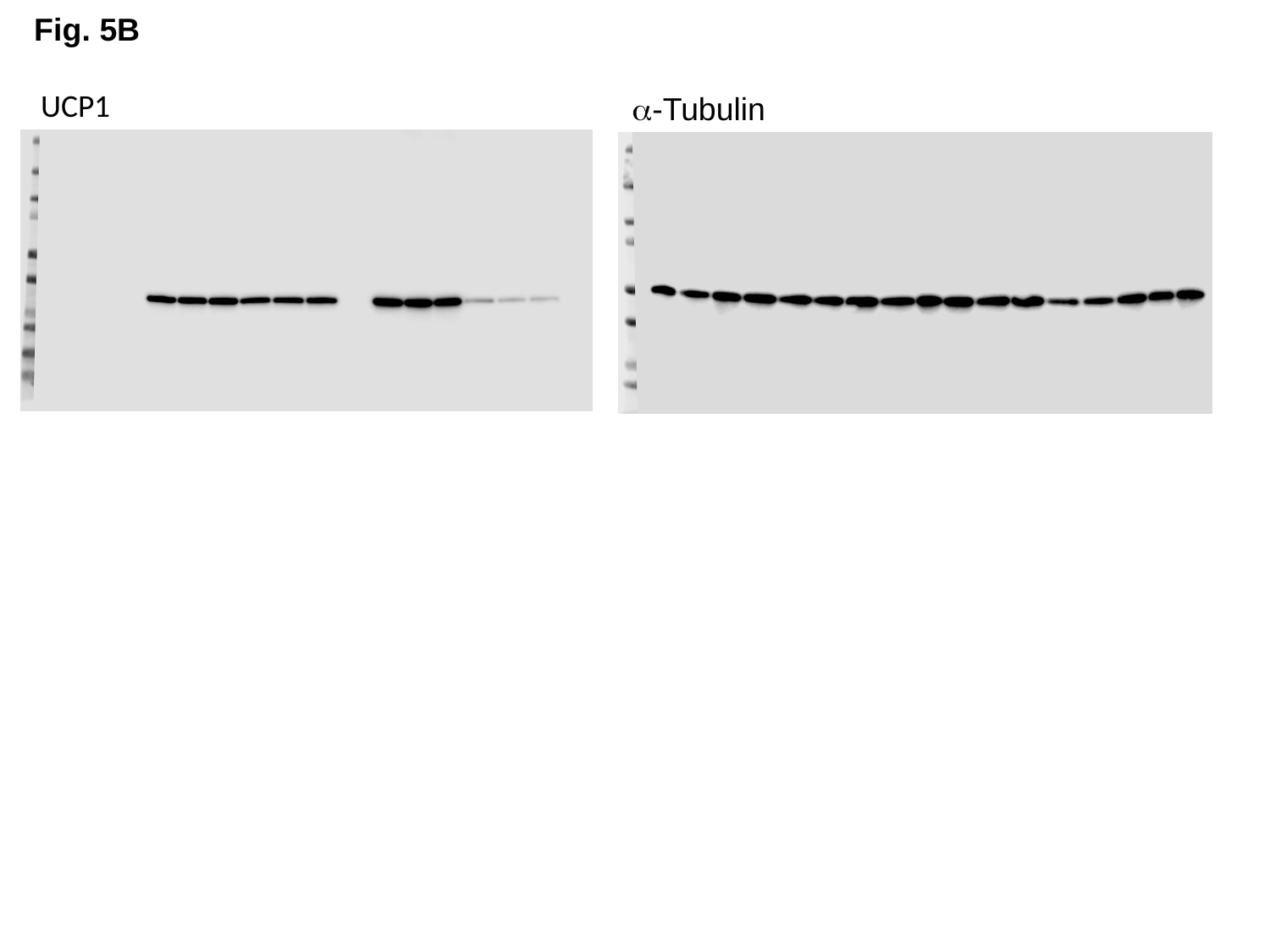

Fig. 5B
UCP1
α-Tubulin

### Slide 14
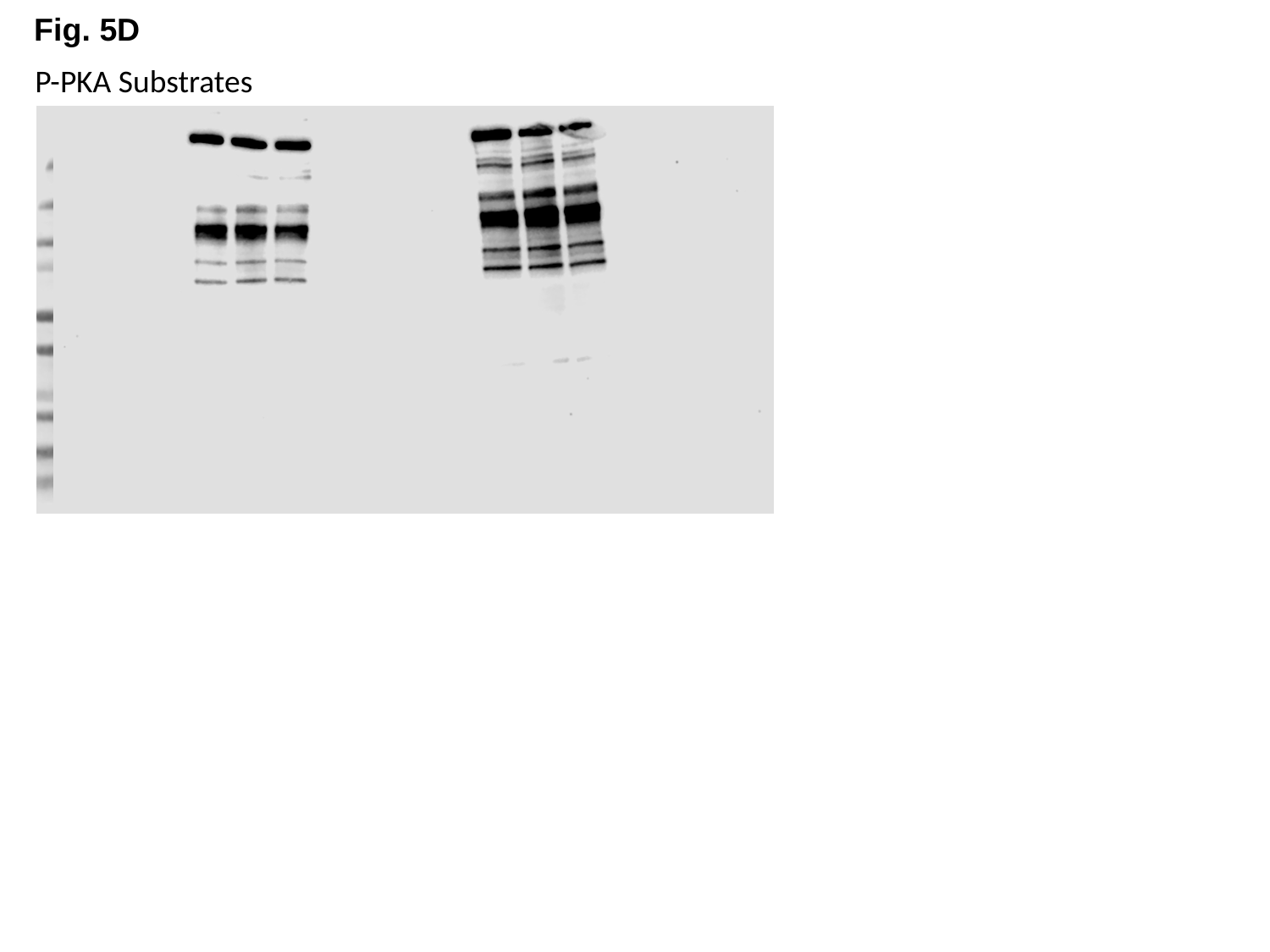

Fig. 5D
P-PKA Substrates

### Slide 15
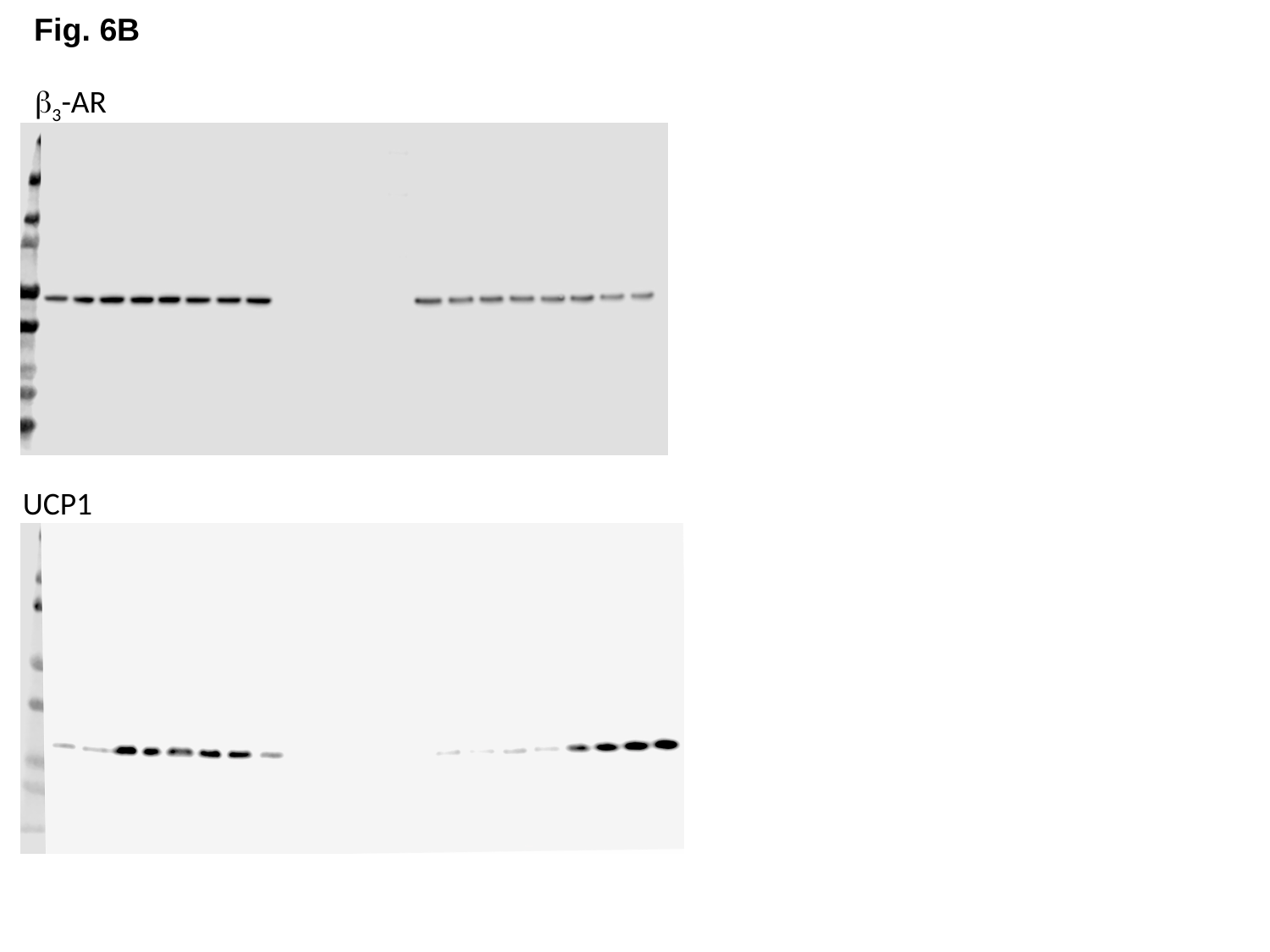

Fig. 6B
β_3_-AR
UCP1

### Slide 16
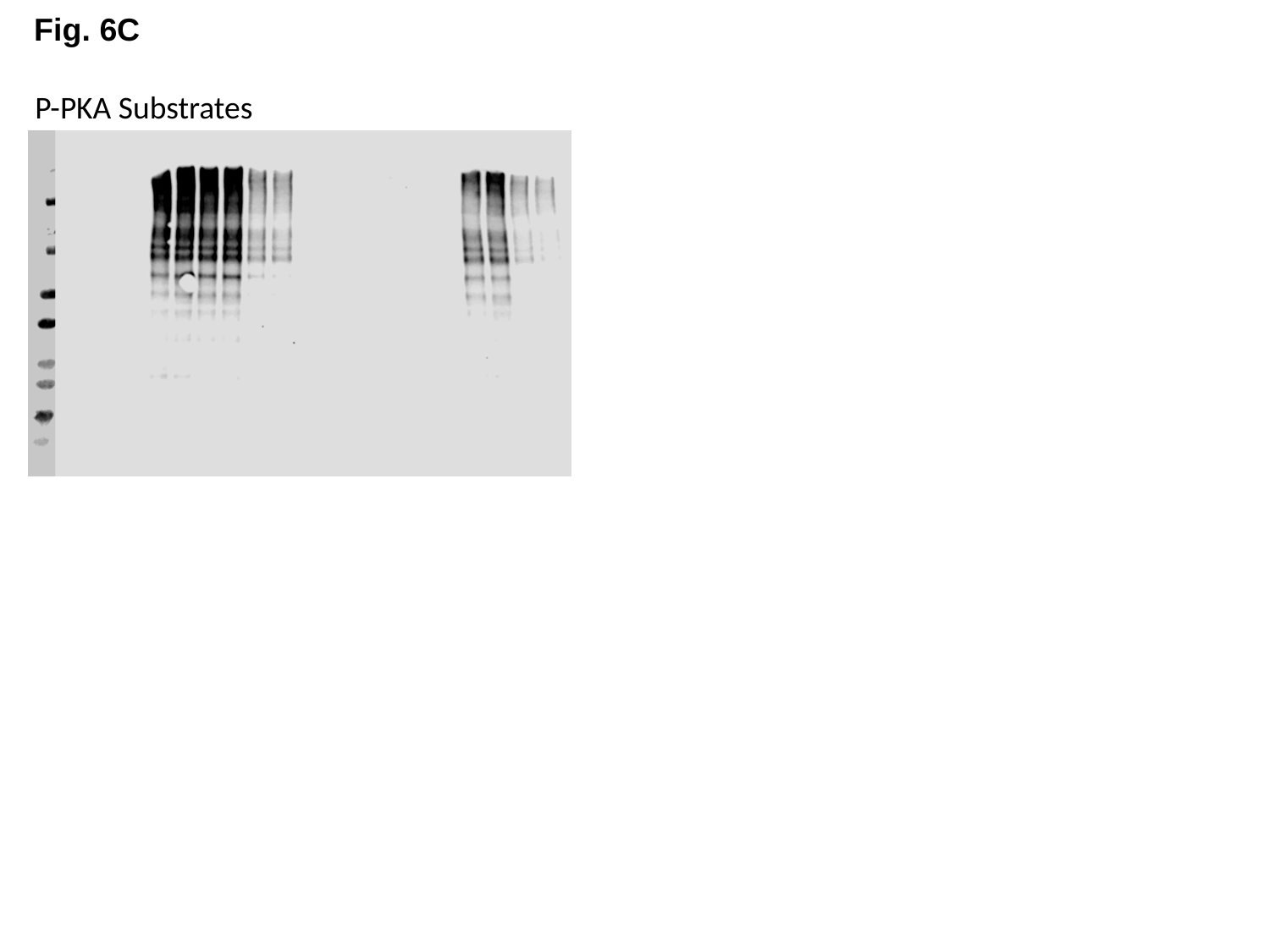

Fig. 6C
P-PKA Substrates

### Slide 17
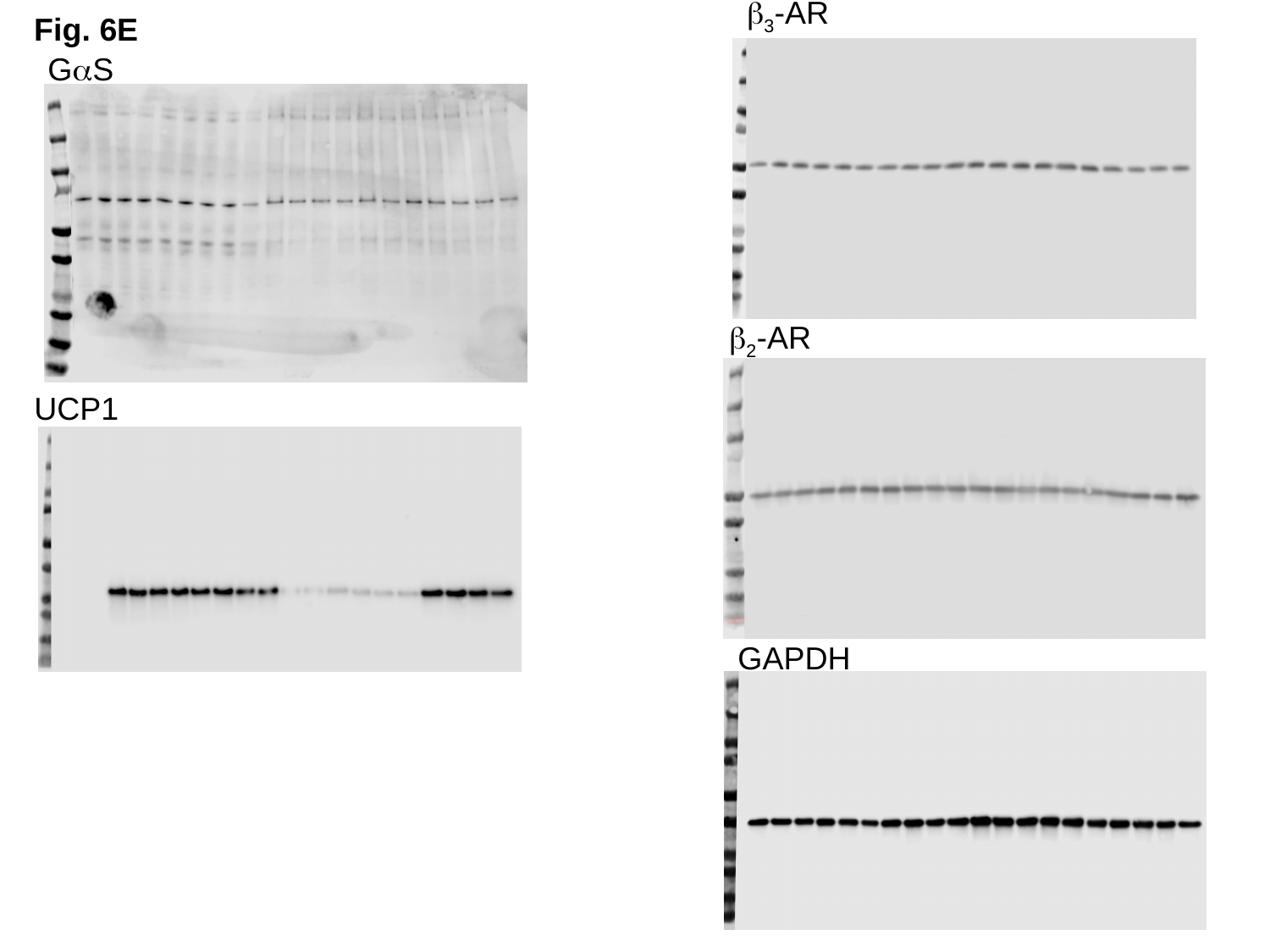

β_3_-AR
Fig. 6E
GαS
β_2_-AR
UCP1
GAPDH

### Slide 18
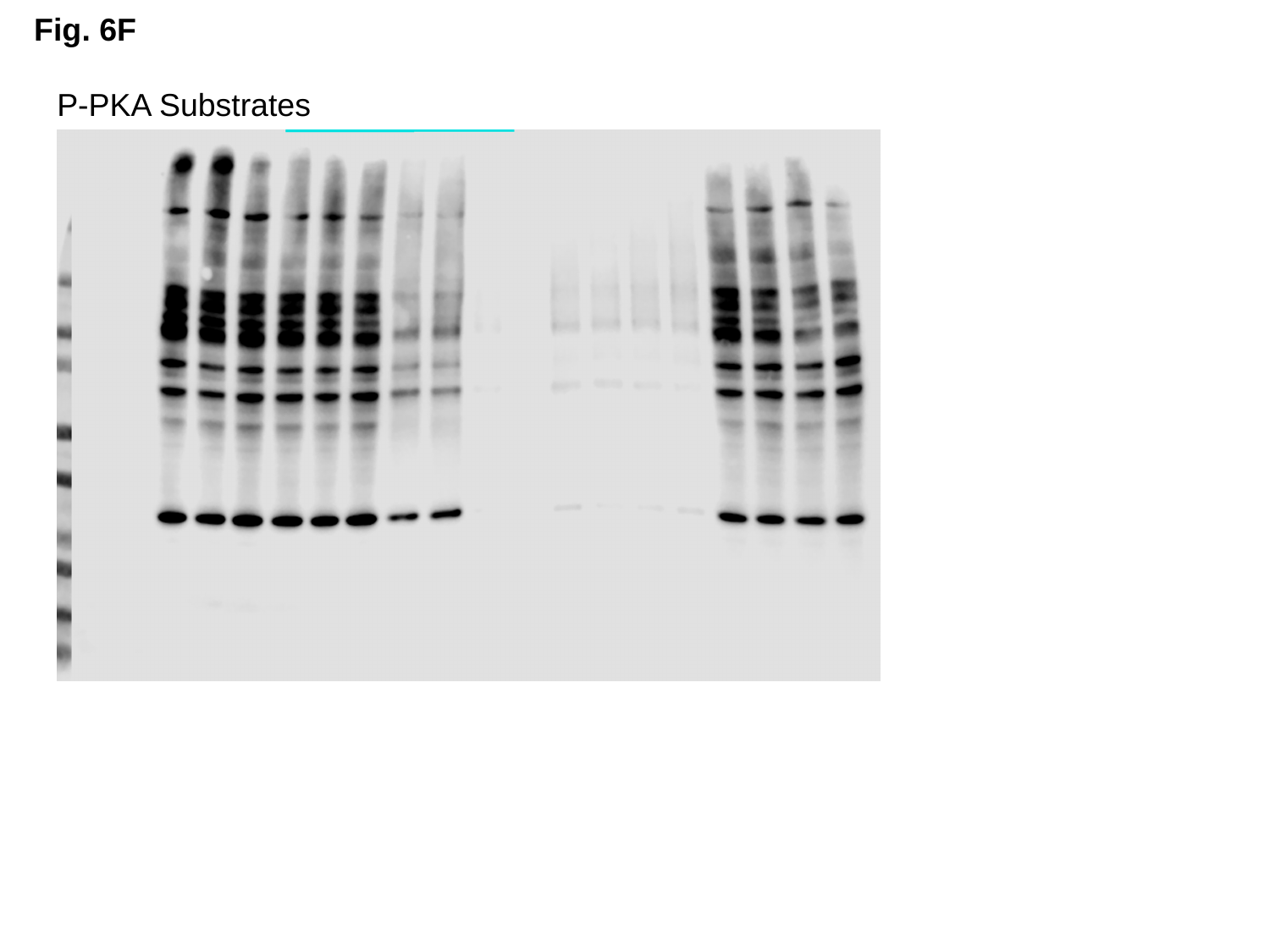

Fig. 6F
P-PKA Substrates

### Slide 19
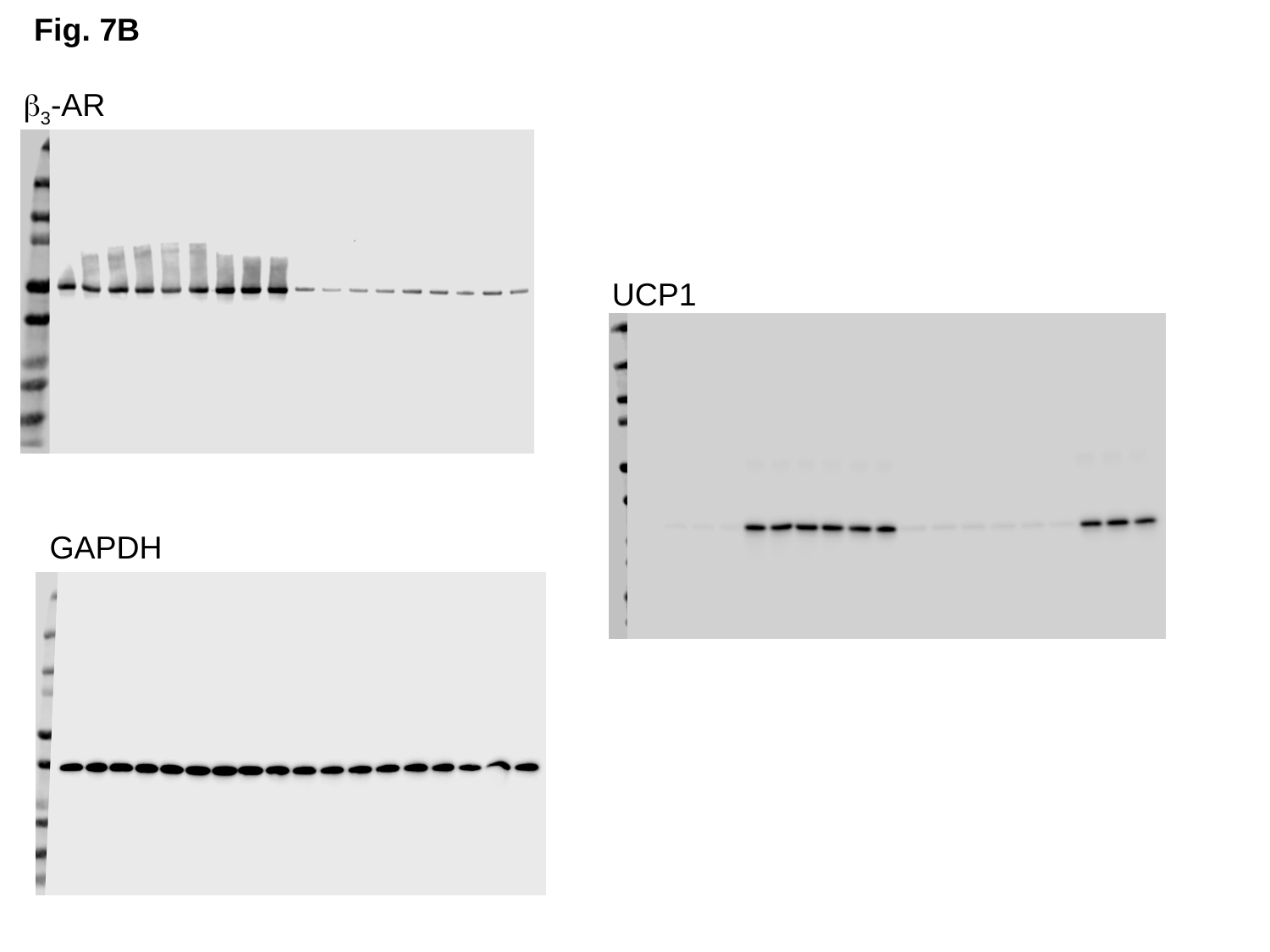

Fig. 7B
β_3_-AR
UCP1
GAPDH

### Slide 20
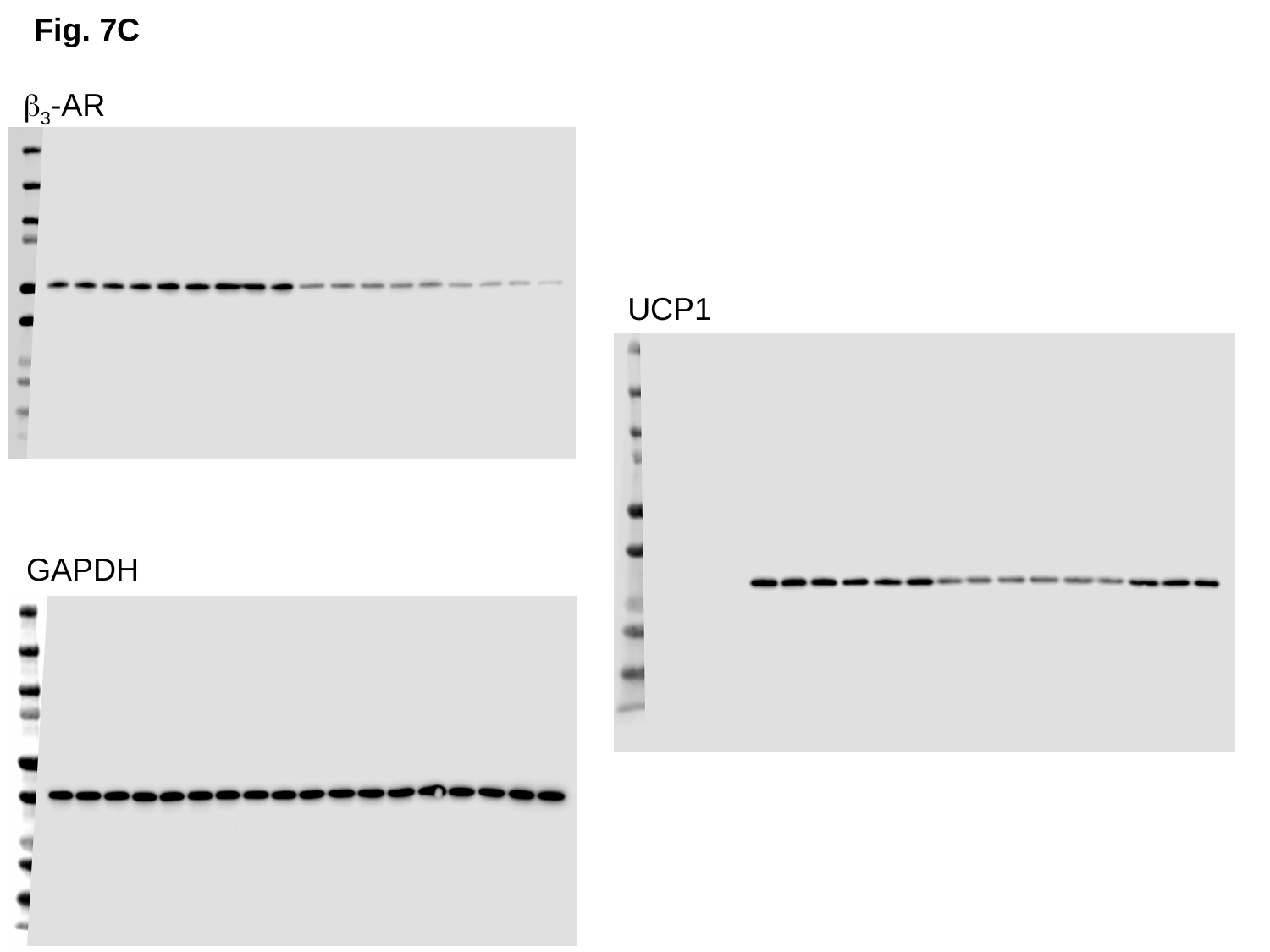

Fig. 7C
β_3_-AR
UCP1
GAPDH

### Slide 21
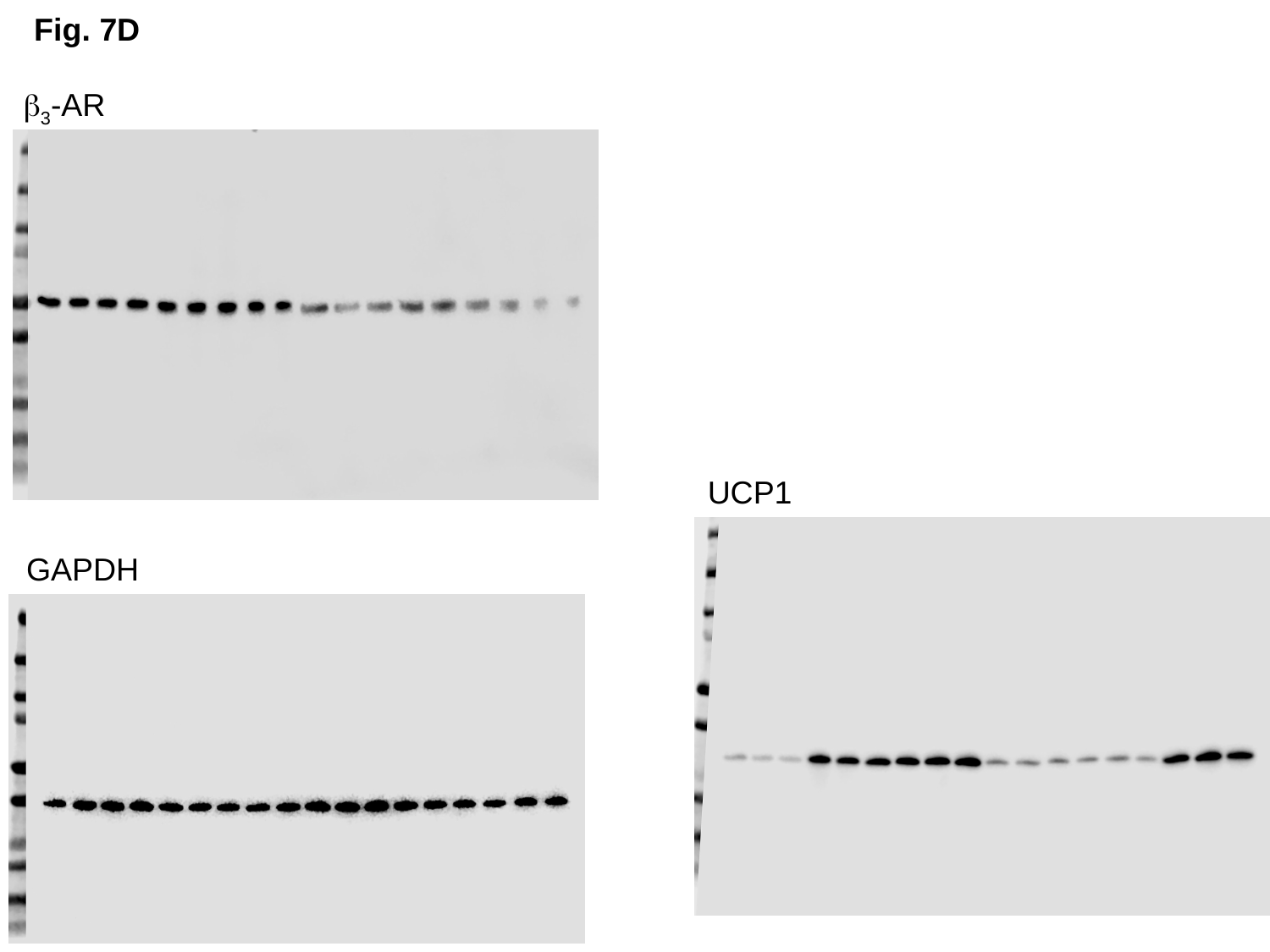

Fig. 7D
β_3_-AR
UCP1
GAPDH

### Slide 22
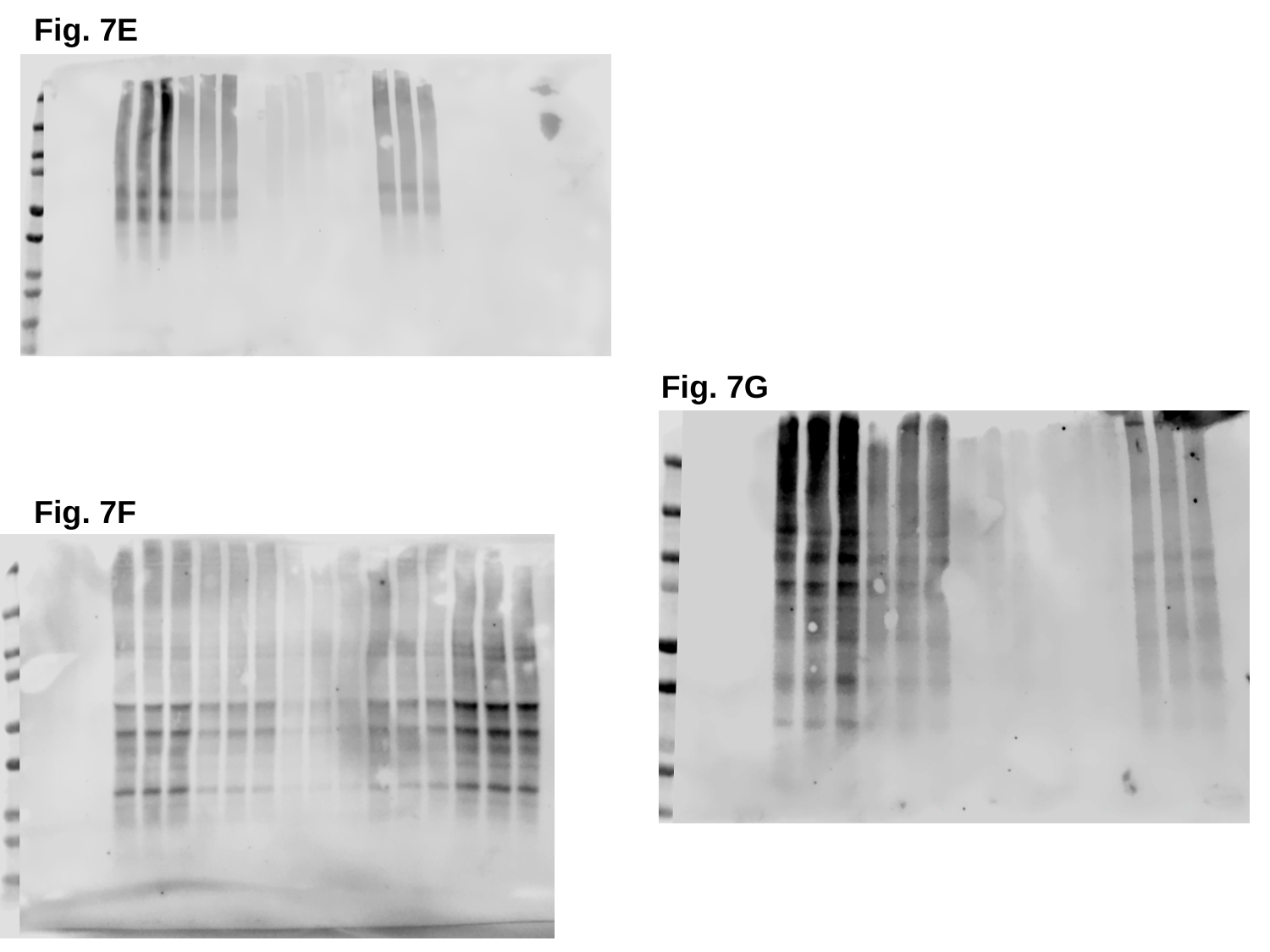

Fig. 7E
Fig. 7G
Fig. 7F

### Slide 23
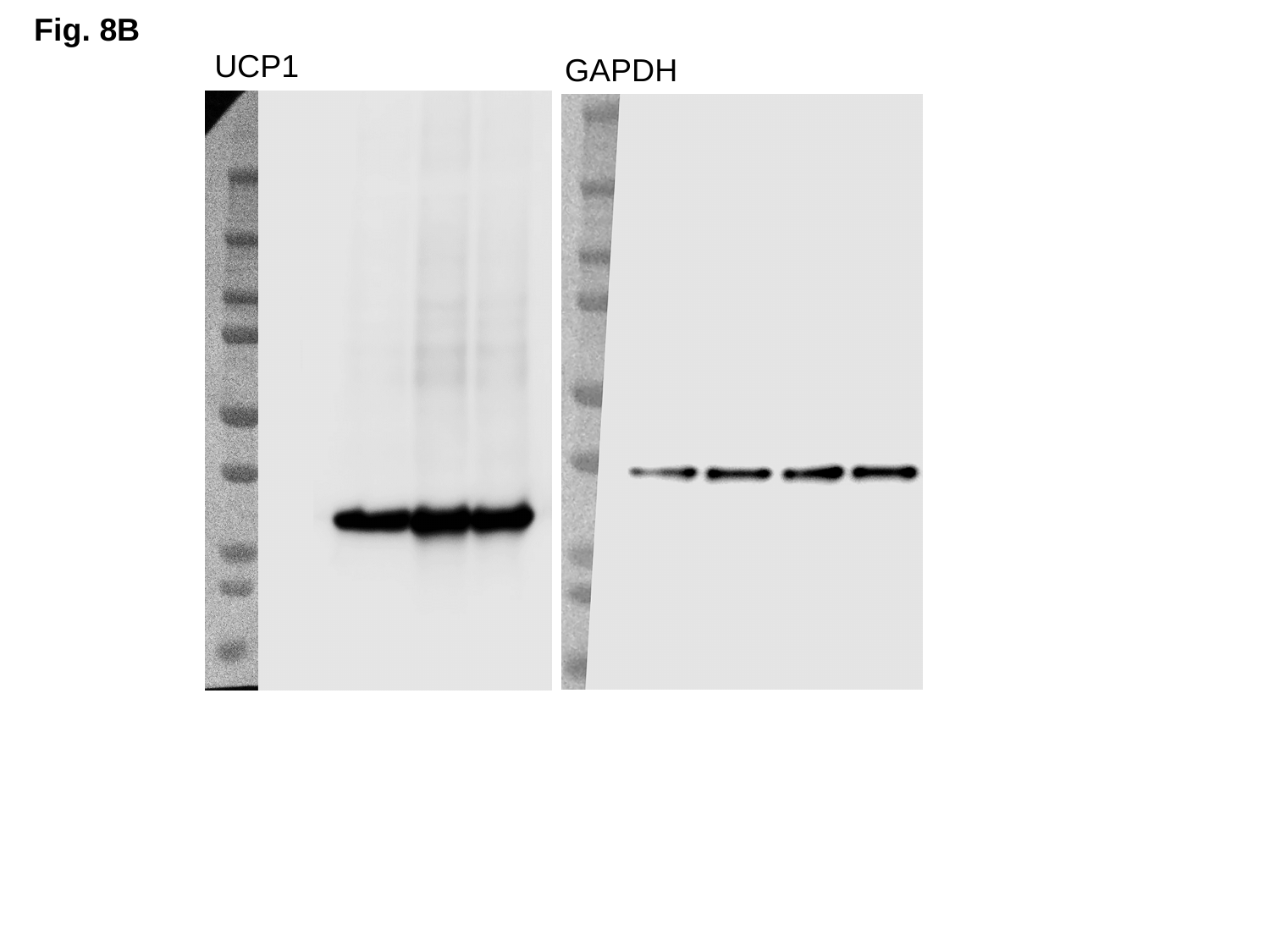

Fig. 8B
UCP1
GAPDH

### Slide 24
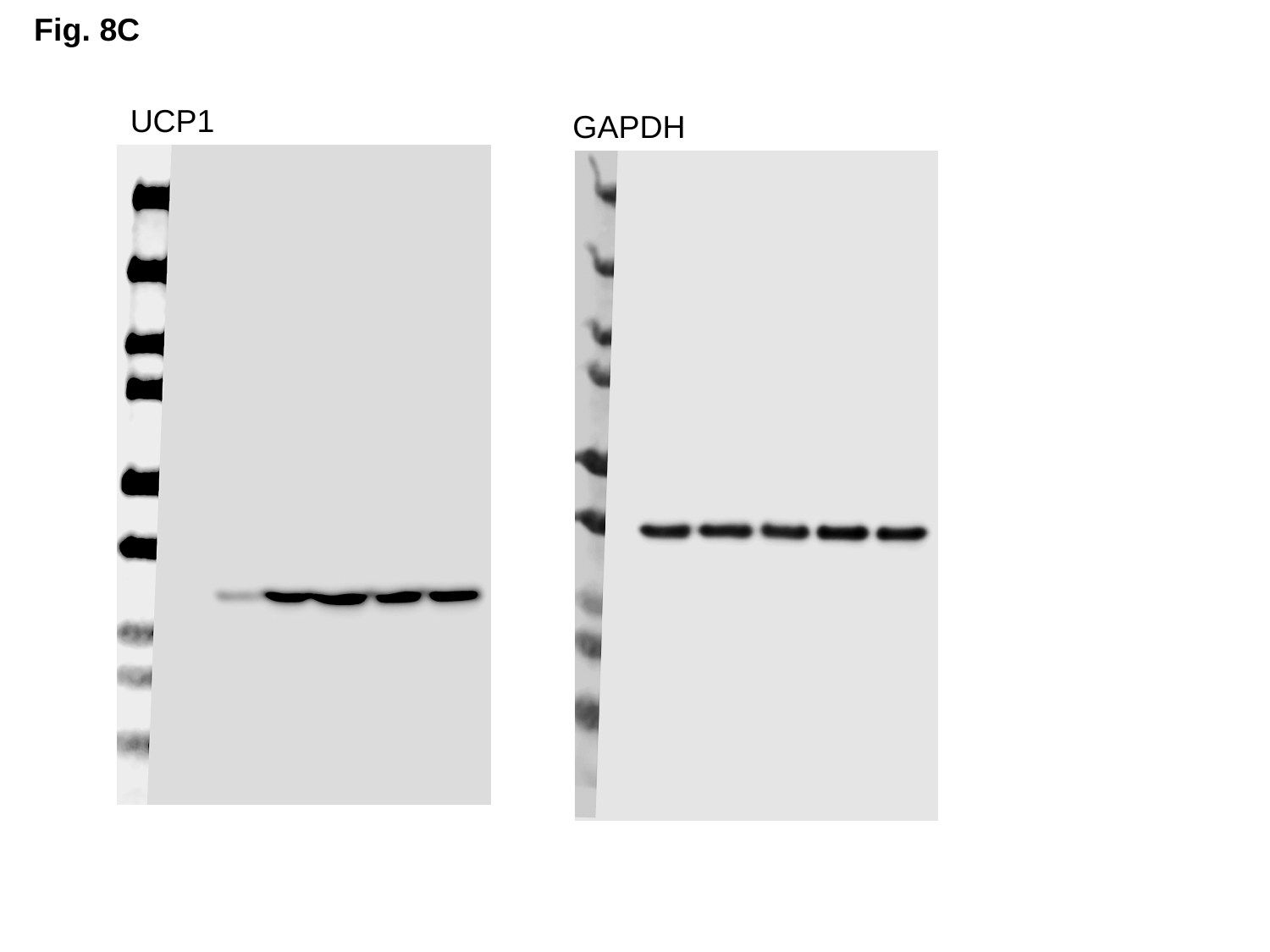

Fig. 8C
UCP1
GAPDH

### Slide 25
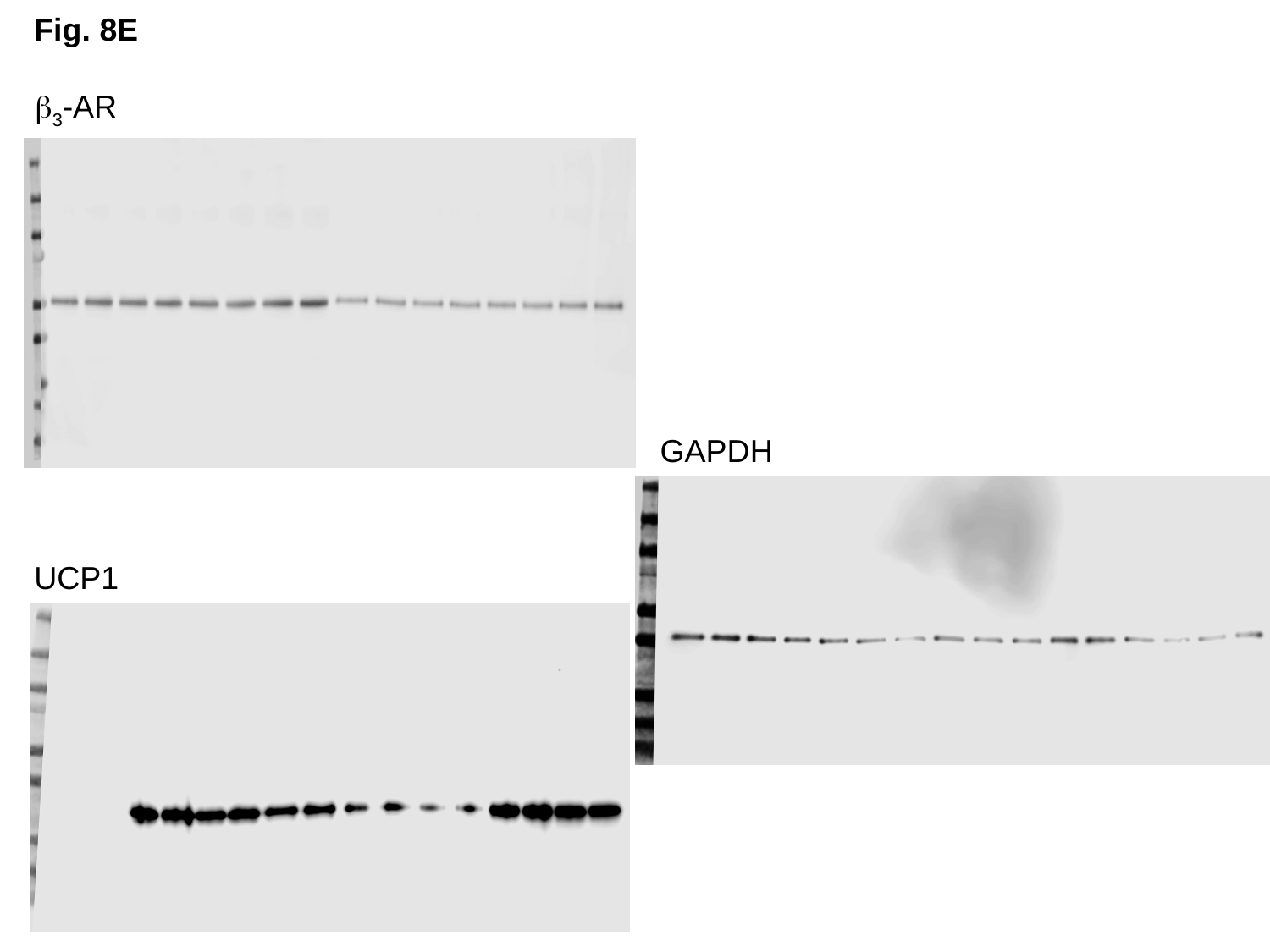

Fig. 8E
β_3_-AR
GAPDH
UCP1

### Slide 26
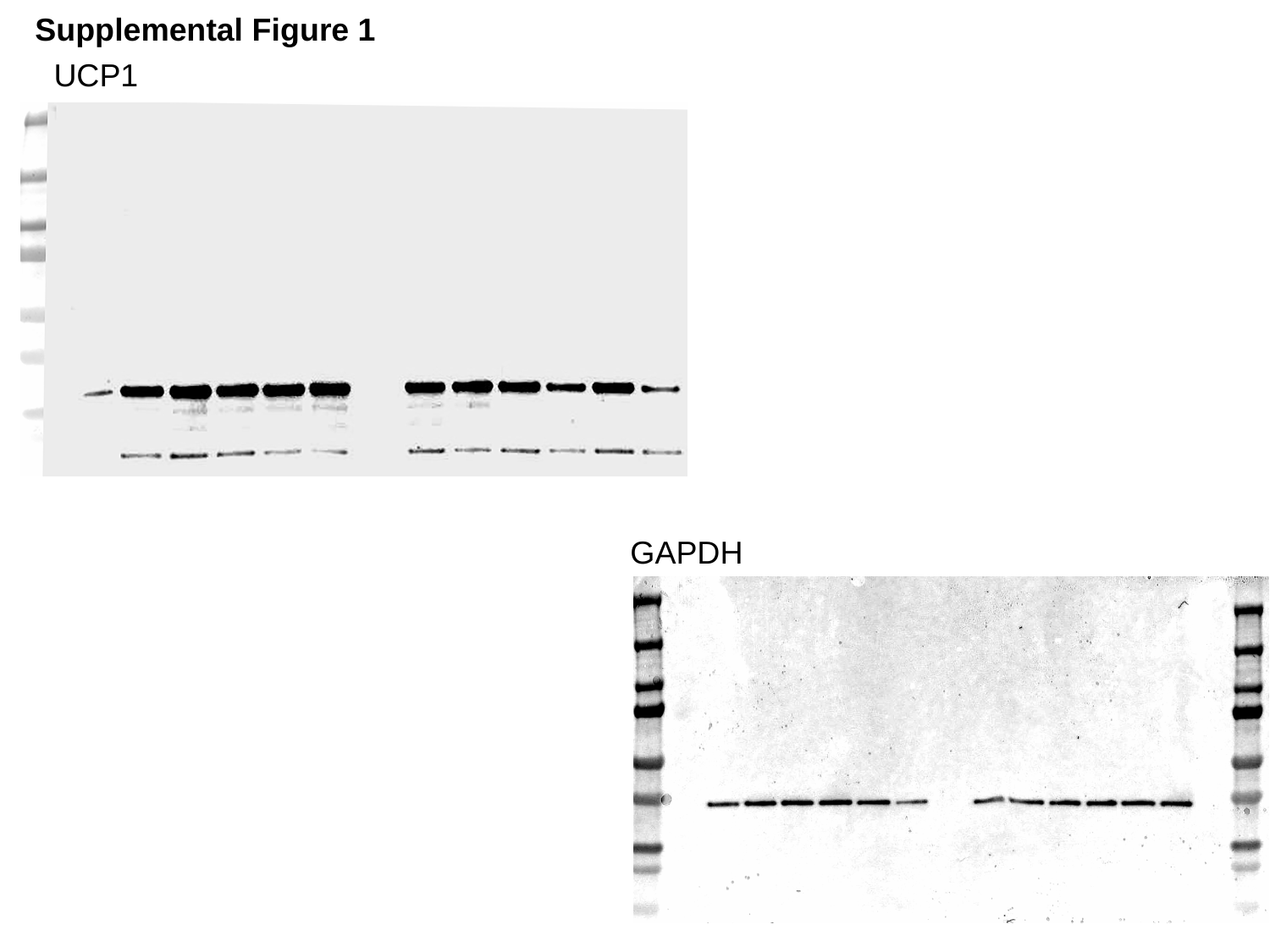

Supplemental Figure 1
UCP1
GAPDH

### Slide 27
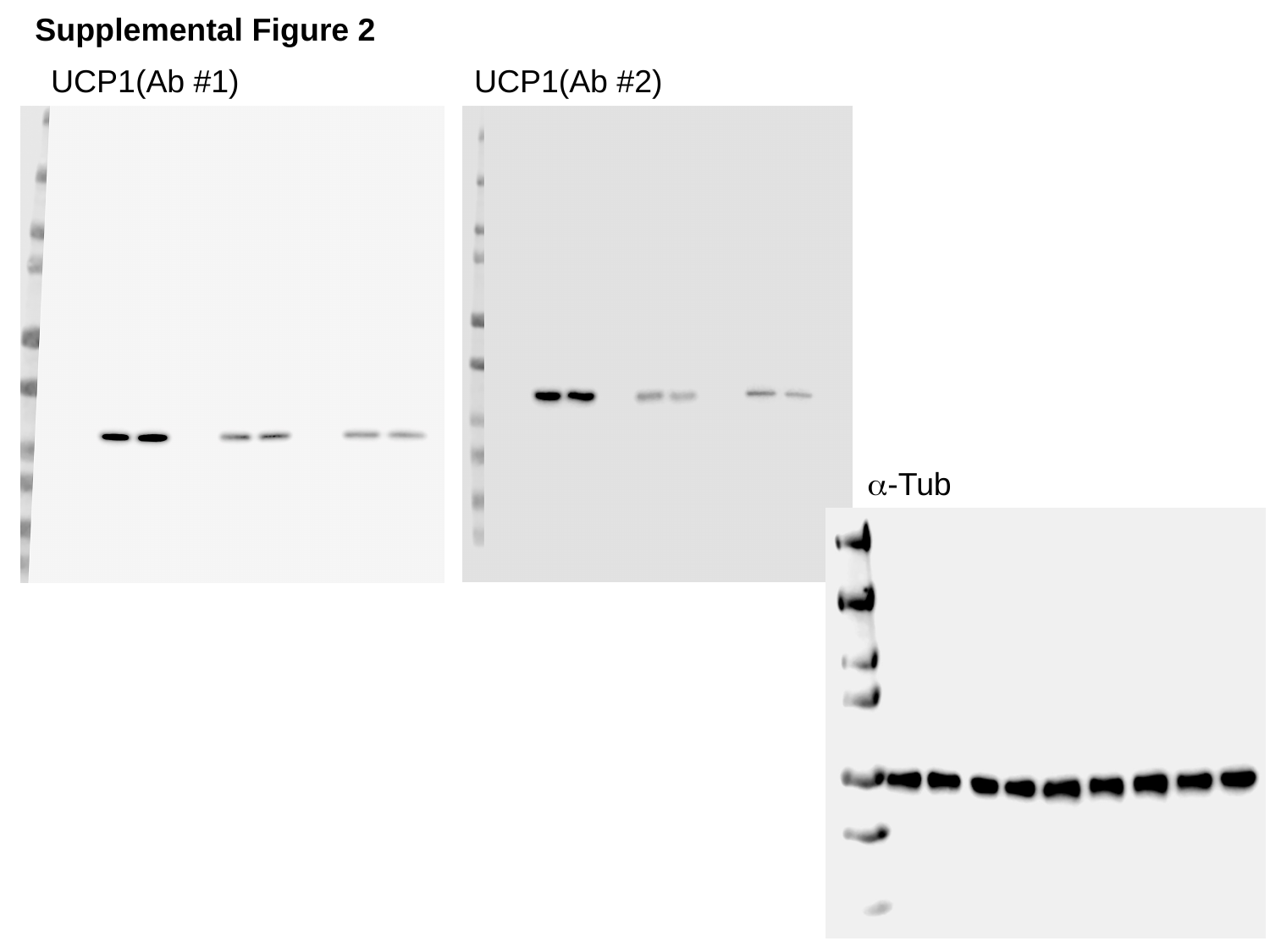

Supplemental Figure 2
UCP1(Ab #1)
UCP1(Ab #2)
α-Tub
